# The transcription factor Pdr802 regulates Titan cell formation, quorum sensing, and pathogenicity of *Cryptococcus neoformans*

**DOI:** 10.1101/2020.06.29.179242

**Authors:** Julia C. V. Reuwsaat, Daniel P. Agustinho, Heryk Motta, Holly Brown, Andrew L. Chang, Michael R. Brent, Livia Kmetzsch, Tamara L. Doering

## Abstract

*Cryptococcus neoformans* is a ubiquitous, opportunistic fungal pathogen that kills almost 200,000 people worldwide each year. It is acquired when mammalian hosts inhale the infectious propagules; these are deposited in the lung and, in the context of immunocompromise, may disseminate to the brain and cause lethal meningoencephalitis. Once inside the host, *C. neoformans* undergoes a variety of adaptive processes, including secretion of virulence factors, expansion of a polysaccharide capsule that impedes phagocytosis, and the production of giant (Titan) cells. The transcription factor Pdr802 is one regulator of these responses to the host environment. Expression of the corresponding gene is highly induced under host-like conditions *in vitro* and is critical for *C. neoformans* dissemination and virulence in a mouse model of infection. Direct targets of Pdr802 include the quorum sensing proteins Pqp1, Opt1 and Liv3; the transcription factors Stb4, Zfc3 and Bzp4, which regulate cryptococcal brain infectivity and capsule thickness; the calcineurin targets Had1 and Crz1, important for cell wall remodeling and *C. neoformans* virulence; and additional genes related to resistance to host temperature and oxidative stress, and to urease activity. Notably, cryptococci engineered to lack Pdr802 showed a dramatic increase in Titan cells, which are not phagocytosed and have diminished ability to directly cross biological barriers. This explains the limited dissemination of *pdr802* mutant cells to the central nervous system and the consequently reduced virulence of this strain. The role of Pdr802 as a negative regulator of Titan cell formation is thus critical for cryptococcal pathogenicity.

**IMPORTANCE:** The pathogenic yeast *Cryptococcus neoformans* presents a worldwide threat to human health, especially in the context of immunocompromise, and current antifungal therapy is hindered by cost, limited availability, and inadequate efficacy. After the infectious particle is inhaled, *C. neoformans* initiates a complex transcriptional program that integrates cellular responses and enables adaptation to the host lung environment. Here we describe the role of the transcription factor Pdr802 in the response to host conditions and its impact on *C. neoformans* virulence. We identified direct targets of Pdr802 and also discovered that it regulates cellular features that influence movement of this pathogen from the lung to the brain, where it causes fatal disease. These findings advance our understanding of a serious disease.

## INTRODUCTION

Cryptococcosis is a fungal infection caused by *Cryptococcus neoformans* and *Cryptococcus gattii. C. neoformans* is a ubiquitous opportunistic pathogen that infects mainly immunocompromised patients, while *C. gattii* is capable of infecting immunocompetent individuals (1). Cryptococcosis causes 180,000 deaths worldwide each year, including roughly 15% of all AIDS-related deaths (2), and is initiated by the inhalation of spores or desiccated yeast cells. In immunocompetent individuals, this typically leads to an asymptomatic pulmonary infection that is controlled by the host immune response, although a population of *C. neoformans* may remain latent for extended periods of time (3–5).

Under conditions of immunocompromise, cryptococci disseminate from the lung to the brain. Mechanisms that have been suggested to mediate fungal crossing of the blood-brain barrier (BBB) include transcellular migration, in which the yeast cells enter and exit vascular endothelial cells (6–9); paracellular movement, in which they cross the BBB at junctions between endothelial cells (10–12); and ‘Trojan horse’ crossing, whereby macrophages harboring *C. neoformans* enter the brain (13). Cryptococcal meningoencephalitis is difficult to treat and frequently lethal, for reasons that include the availability and cost of therapy (14, 15).

The ability of *C. neoformans* to survive and proliferate in the lung, and subsequently disseminate to the brain, depends on viability at mammalian body temperature and the expression of multiple virulence traits; these include secreted factors (16, 17), a polysaccharide capsule that surrounds the cell wall (18), and the production of giant (Titan) cells (19, 20). One secreted molecule, the pigment melanin, associates with the cell wall, where its antioxidant properties protect fungal cells from reactive oxygen species produced as a host immune defense (21–25). Urease, a secreted metalloenzyme that converts urea to ammonia and CO_2_, may affect the course of infection by modulating environmental pH and damaging host tissue structure (11, 12, 26).

The capsule, composed primarily of large polysaccharides (27–29), is a key cryptococcal virulence factor that impairs phagocytosis by immune cells (30–35). This dynamic entity changes its size and structure during interactions with the host or external environment (36–39), contributing to fungal adaptation (40, 41). Capsule polysaccharides that are shed from the cell enable diagnosis of cryptococcal infection and also impede host responses (35, 42).

Titan cells display a cryptococcal morphotype that has been variously characterized as having cell body diameter (excluding the capsule) greater than 10 or 15 µm or total cell diameter (including the capsule) that exceeds 30 µm (20, 43, 44). These cells are polyploid and produce normal-size cells during infection (19, 45, 46). Titan cell formation is triggered by exposure to the host environment, including nutrient starvation, reduced pH, and hypoxia (47–49), although the extent of induction depends on the host immune response and the duration of infection (45, 50). Titan cell production appears to benefit the development of pulmonary *C. neoformans* infection, since these large cells are less susceptible to internalization by host phagocytes and more resistant to oxidative stress than normal-size cells (19, 46). Some of these effects may be explained by the highly cross-linked capsule and thickened cell wall of Titan cells (51). In contrast to their success in the lungs, Titan cells show impaired dissemination to the brain (19, 46).

*C. neoformans* experiences a dramatic change in conditions upon entering a host, including altered nutrient levels and pH. To adapt to the new environment, cryptococci activate a network of transcription factors (TFs) (39, 52). For example, imbalances in ion homeostasis trigger transcriptional changes mediated by the TFs Zap1 (53), Cuf1 (54), Pho4 (55), Cir1 (56), and Crz1 (57). Alkaline pH stimulates expression of the TF Rim101, which enables growth under basic conditions and other stresses such as high salt and iron limitation; it also promotes the association of capsule polysaccharide with the cell and the formation of Titan cells (47, 58).

Overlapping TF circuits regulate cryptococcal virulence determinants, including polysaccharide capsule production and melanin synthesis. For example, Usv101, an important regulator of capsule thickness and polysaccharide shedding, also regulates three other TFs (Gat201, Crz1, and Rim101) and multiple polysaccharide-related enzymes (59). Gat201 further regulates additional virulence-related transcription factors and the anti-phagocytic protein Blp1 (60), while Crz1 plays a central role in the maintenance of plasma membrane and cell wall stability (57, 61, 62). Crz1 expression is also modulated by the calcineurin signaling pathway, which is required for normal yeast growth at 37°C, virulence, and sexual reproduction (63). A group of TFs, including Usv101, Bzp4, Hob1, and Mbs1 (59, 64), act together to regulate melanin production; deletion of Bzp4 also alters capsule (52).

In this study, we investigated the TF Pdr802. The corresponding gene has a high rate of non-synonymous mutations, which suggests it is evolving rapidly (65). Pdr802 has previously been implicated in *C. neoformans* virulence (39, 52, 66), but its specific role and targets are not known. We discovered that Pdr802 is induced in host-like conditions, is a negative regulator of Titan cell formation, and influences capsule thickness and phagocytosis by macrophages. It also regulates genes whose products act in cell wall remodeling, virulence factor production, resistance to host temperature and oxidative stress, and quorum sensing. These functions make Pdr802 critical for cryptococcal survival in the lung and dissemination to the brain.

## RESULTS

### The role of Pdr802 in *C. neoformans* virulence

The importance of Pdr802 in *C. neoformans* virulence has been demonstrated in multiple experimental models. Liu and collaborators first reported in 2008 that partial deletion of *PDR802* reduced *C. neoformans* infectivity in a competition assay of pooled *C. neoformans* strains (66). In 2015, Maier *et al* showed that a *pdr802* deletion mutant had reduced virulence when tested individually in a short-term mouse model of infection (39). Later that year, Jung and colleagues reported that Pdr802 was required for full virulence in both wax moth larvae and short-term mouse infection using pooled strains (52). Most recently, Lee and collaborators showed that Pdr802 was required for brain infection (67).

To further investigate the role of Pdr802 in pathogenesis, we complemented a complete deletion strain in the KN99α background that we had previously generated (*pdr802*) (39) with the intact gene at its native locus (*PDR802*). To examine targets of Pdr802, we also constructed a strain that expresses the protein fused to mCherry at its N-terminus (Figure S1A). All of these strains lacked or expressed RNA encoding *PDR802* or its modified forms as expected (Figure S1B) and *PDR802* was expressed at wild-type levels in the complemented and modified strains (Figure S1C).

We next assessed the long-term survival of C57BL/6 mice infected with the parental wild-type strain (KN99α), the deletion mutant (*pdr802*), or the complemented mutant (*PDR802*). In this model, mice infected with the parent or complemented strains survived for roughly three weeks, while those infected with the deletion mutant showed a striking increase in survival: all animals survived for at least 65 days and over half survived to the end of the study (100 days; Figure 1A). The lung burden measured at the time of death for *pdr802*-infected mice in this study was approximately 100-fold lower than that of wild type infections (Figure S2A), demonstrating the importance of this TF in *C. neoformans* virulence. Mean brain burden at the time of death was more similar between mutant and wild type infections (Figure S2A), although we did note some heterogeneity in this measure for *pdr802*-infected mice; animals sacrificed at around two months of infection (red symbols) showed brain burden similar to WT levels, while brain burden of mice sacrificed at day 100 (blue symbols) ranged between zero fungal cells and WT level.

**Figure 1.**
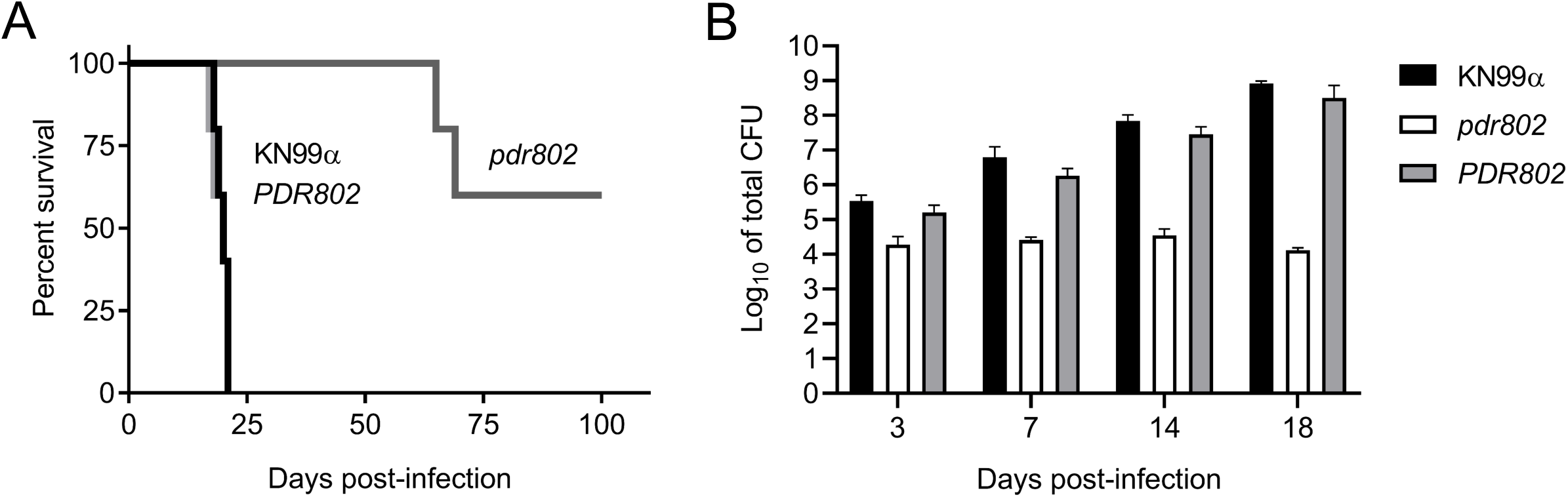
The transcription factor Pdr802 influences *C. neoformans* virulence. A. Survival of C57BL/6 mice over time after intranasal inoculation with 5 x 10^4^ cryptococci of the strains indicated, with sacrifice triggered by weight below 80% of peak. B. Mean +/− SD of total colony-forming units (CFU) in lung tissue at various times post-infection. CFU inoculated for each strain were 41,400 (KN99α), 42,800 (*pdr802*) and 26,600 (*PDR802*). p<0.05 for *pdr802* compared to the other strains at all time points.

We next examined the time course of fungal proliferation in the lungs. As expected, the burdens of WT and the complemented mutant strains increased steadily over an 18-day interval (Figure 1B), eventually reaching roughly 10^5^ times the original inoculum. Towards the end of this period, these cells were also detected in the blood and brain (Figure S2B). In contrast, the lung burden of *pdr802* remained close to the inoculum throughout this period, with no mutant cells detected in the blood or brain. At a late time point of *pdr802* infection (75 days), we again noted some heterogeneity of fungal burden: one mouse had high lung burden with no dissemination, another had high lung burden with moderate brain burden, and the third had extremely low lung burden with no dissemination (Figure S2C). No colony-forming units (CFU) were detected in the blood of *pdr802*-infected mice at any point during infection. These results suggest that even though the *pdr802* mutant is generally hypovirulent and remains at low levels in the lung, it can occasionally reach the brain and, given enough time, accumulate there (see Discussion).

Given the dramatic effects of Pdr802 on fungal virulence, we wondered about the specific biological processes in which this transcription factor is involved. We first examined the behavior of the *pdr802* strain *in vitro*, including stress conditions that might be encountered in the host. We saw no differences in growth of the mutant compared to WT cells under conditions that challenge cell or cell wall integrity, including the presence of sorbitol, high salt, cell wall dyes, caffeine, sodium dodecyl sulfate (SDS), or ethanol (Figure S3A-C). The mutant also showed no altered susceptibility to elements of the host response, such as nitrosative or oxidative stresses, or in melanin production. All of these results held whether growth was at 30°C, 37°C, or 37°C in the presence of 5% CO_2_, which was recently described as an independent stress for *C. neoformans* (68) (Figure S3A-C). Finally, the mutant showed no difference from wild-type cells in secretion of urease at 30°C or 37°C (Figure S3D).

### Pdr802 is regulated by “host-like” conditions

We next tested the growth of the *pdr802* mutant under conditions more like those encountered inside the mammalian host, using tissue culture medium (DMEM) at 37°C in the presence of 5% CO_2_. We found that although the *pdr802* mutant grew like WT in rich medium (YPD), it grew poorly in DMEM (Figure S4A-B). To test whether the mutant cells were dead or just static after growth in DMEM, we plated aliquots on solid medium to measure CFU over time (Figure 2A). The *pdr802* culture showed a dramatic decrease in viability compared to WT and the complemented strain, which was greatest in the first 24 h. This is the same time frame in which expression of the *PDR802* gene shows a striking increase in wild type cells, as measured by RNA-Seq (Figure 2B).

**Figure 2.**
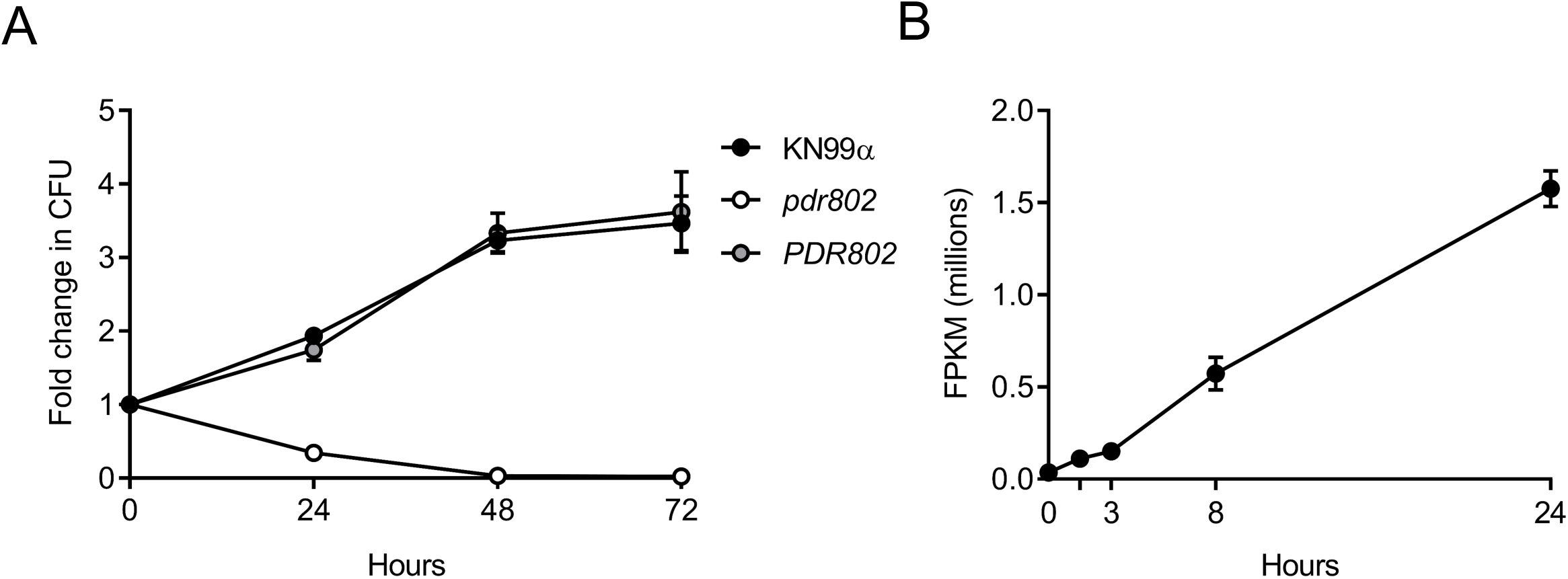
*PDR802* expression is required for cell viability and induced during growth in host-like conditions. A. Cells grown in DMEM at 37°C and 5% CO_2_ were sampled at the times indicated and plated on YPD to assess viability (measured by CFU and plotted as fold-change from time 0). B. *PDR802* expression in KN99α cells grown in DMEM at 37°C and 5% CO_2_ was assessed by RNA-seq as in Li *et al*., 2018 (116).

Another important feature that is induced by growth in DMEM at 37°C and 5% CO_2_ is the polysaccharide capsule, which we previously reported to be regulated by Pdr802, based on negative staining with India ink (39). Fluorescence microscopy confirmed increased capsule thickness of the mutant, which reverted to WT in the complemented strain (Figure 3A). To quantify this change, we took advantage of a semi-automated assay that we have developed (Figure S5), which measures capsules on a population scale (Figure 3B) and is therefore very sensitive. This analysis showed that the capsule thickness of *pdr802* cells resembles that of the well-studied hypercapsular mutant *pkr1* (39, 69, 70) and is completely restored to WT by complementation at the native locus (Figure 3C). Previous studies suggest that capsule thickness upon induction reflects the size of the dominant capsule polymer (glucuronoxylomannan; GXM) (71, 72), which can be analyzed by agarose gel migration and blotting with anti-capsule antibodies (71). Consistent with the difference we observed in capsule thickness by imaging, this method showed decreased mobility of GXM from *pdr802* as capsule induction progressed (Figure S4C).

**Figure 3.**
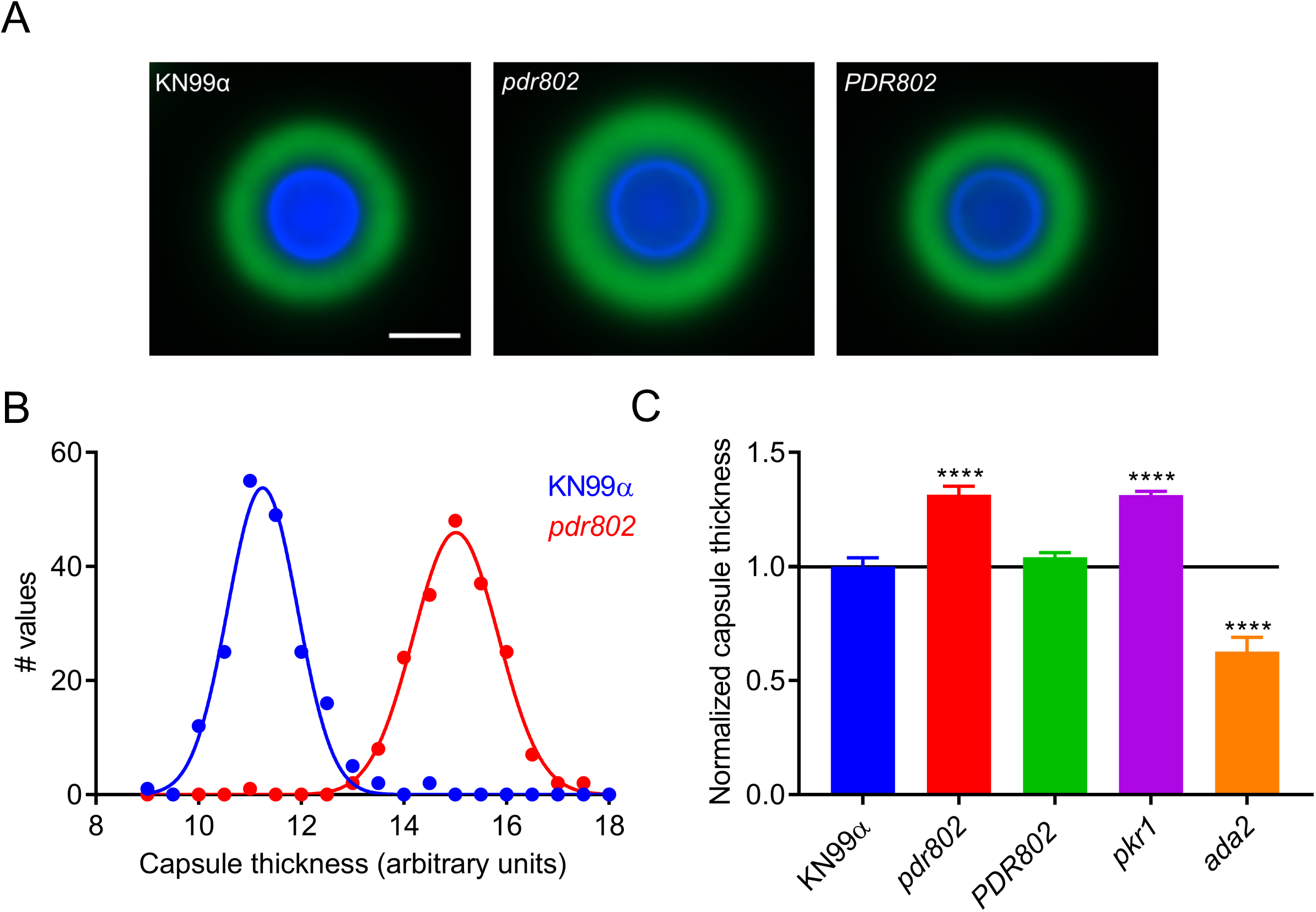
The *pdr802* mutant is hypercapsular. A. Representative immunofluorescence micrographs of the indicated strains after growth in DMEM (37°C, 5% CO_2_) for 24 hours. The capsule was stained with monoclonal antibody anti-GXM 302 conjugated with Alexa 488 (green) and the cell wall with Calcofluor White (blue). All images are to the same scale; scale bar, 5 µm. B. Capsule thickness distribution for the indicated strains. C. Mean +/− SD of capsule size, quantified as detailed in the Methods and Figure S5, with *pkr1* (39) and *ada2* (105) shown as hypercapsular and hypocapsular controls, respectively. ****, p<0.0001 compared to KN99α by one-way ANOVA with posthoc Dunnett test.

To validate the observations that we had made in standard ‘host-like’ conditions based on synthetic tissue culture medium, we conducted similar studies in mouse serum at 37°C and 5% CO_2_. These conditions induced an even more pronounced hypercapsular phenotype of the *pdr802* mutant (Figures 4A and 4B), as well as reduced cell viability (Figure 4C) and increased cell body diameter (Figure 4D).

**Figure 4.**
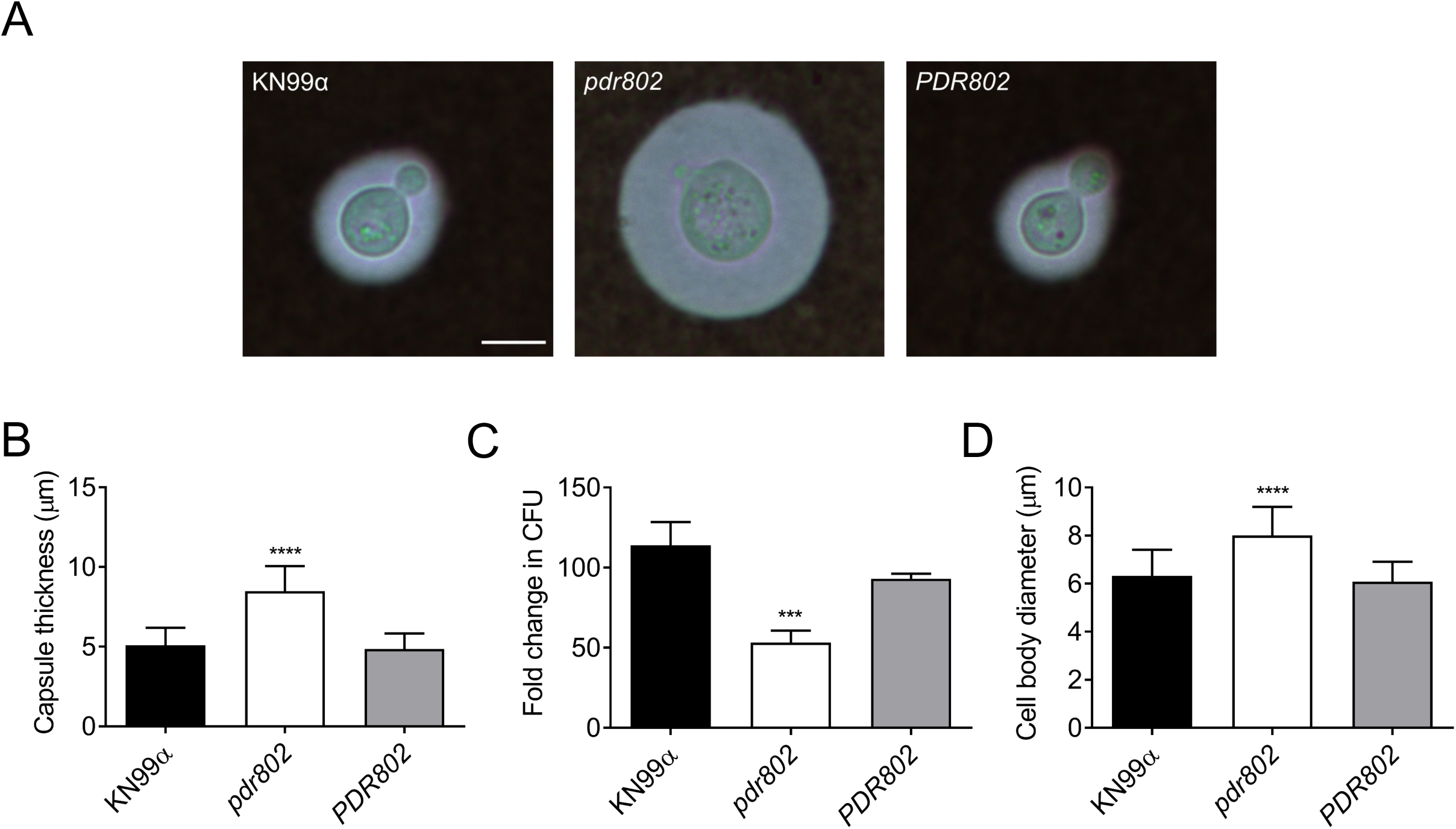
Growth in mouse serum elicits increased capsule thickness and cell body diameter in the *pdr802* mutant. A. Light micrographs of the indicated strains after growth in mouse serum (at 37°C, 5% CO_2_) for 24 h and negative staining with India ink to visualize the capsule. All images are to the same scale; scale bar, 5 µm. B. Mean +/− SD of capsule thickness, assessed by measuring at least 50 cells per strain with ImageJ. C. Cells grown as in Panel A were plated on YPD to assess CFU. Mean +/− SD of the fold-change compared to 0 h is shown. D. Mean +/− SD of cell body diameter, measured as in B. ***, p<0.001 and ****, p<0.0001 for comparison of *pdr802* results to KN99α by one-way ANOVA with posthoc Dunnett test.

We were intrigued by the enlarged cell body and capsule of the *pdr802* mutant cells in host-like conditions *in vitro* and decided to examine these phenotypes *in vivo*. For these studies, we isolated fungal cells from the lungs of mice at various times after infection and assessed their morphology by negative staining (Figure 5A). At each time point, the mean mutant cell body diameter was larger than that of the controls. Additionally, while this parameter was stable for WT and complemented strains throughout the infection period, it trended larger at the end of the infection period for the deletion mutant (Figure 5B). In contrast, mutant capsule thickness, although initially larger than that of control cells, changed little throughout the period, while capsule thickness of control cells increased to that level or beyond (Figure 5C). Furthermore, although the total cell diameter of *pdr802* cells consistently exceeded that of WT and complemented cells, their sizes became more comparable late in infection (Figure S6A). Over time, therefore, the ratio of total cell diameter to cell body diameter for WT and *PDR802* cells steadily increased, while it remained roughly constant for the mutant (Figure S6B).

**Figure 5.**
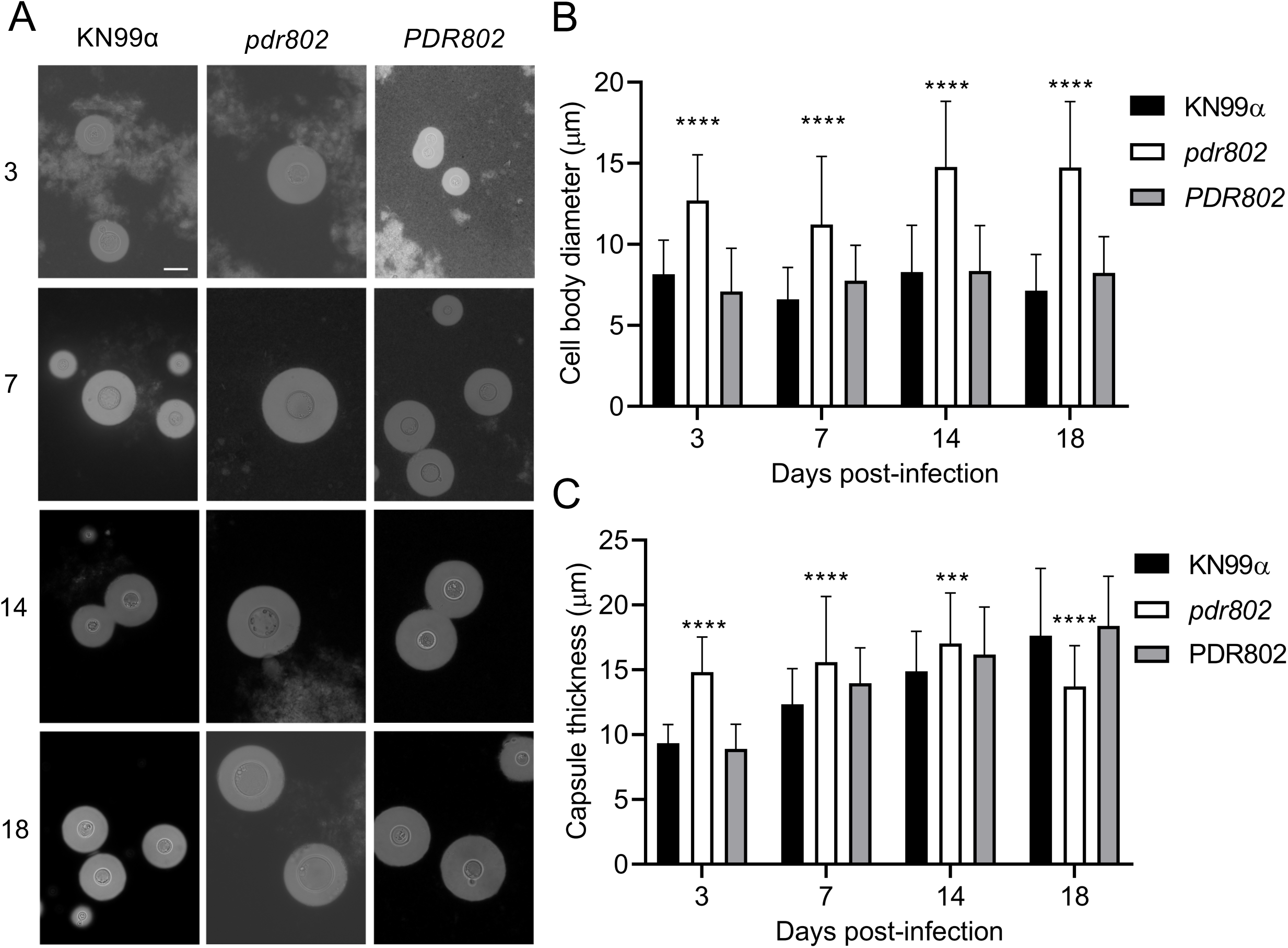
Absence of *PDR802* yields enlarged cells and loss of capsule induction in the context of animal infection. A. India ink staining of fungi isolated from the lungs of mice infected with the indicated strains. Numbers at left indicate the days post-infection. All images are to the same scale; scale bar, 10 µm. B and C. Mean +/− SD of cell body diameter (B) and capsule thickness (C), assessed by measuring at least 50 cells per strain with ImageJ. ****, p<0.0001 and ***, p<0.001 for comparison of *pdr802* results to KN99α or *PDR802* by one-way ANOVA with posthoc Dunnett test for each day post-infection.

### Pdr802 negatively regulates Titan cell formation

We were particularly interested in the cell size phenotype of *pdr802* because Titan cells have been strongly implicated in cryptococcal pathogenesis (19). By any definition of this morphotype (cell body diameter greater than 10 or 15 µm or total cell diameter greater than 30 µm), our mutant cell populations were dramatically enriched in Titan cells at every time of infection that we assessed (Figure S6C).

To specifically test Titan cell formation by the *pdr802* strain, we subjected mutant cells to *in vitro* conditions that induce this process (49) and analyzed the resulting population by flow cytometry. Consistent with our *in vivo* observations, Titan cells constituted a much larger fraction of the population in the mutant culture (13.2%) than in the WT and complemented cultures (1.62% and 1.40%, respectively) (Figure 6).

**Figure 6.**
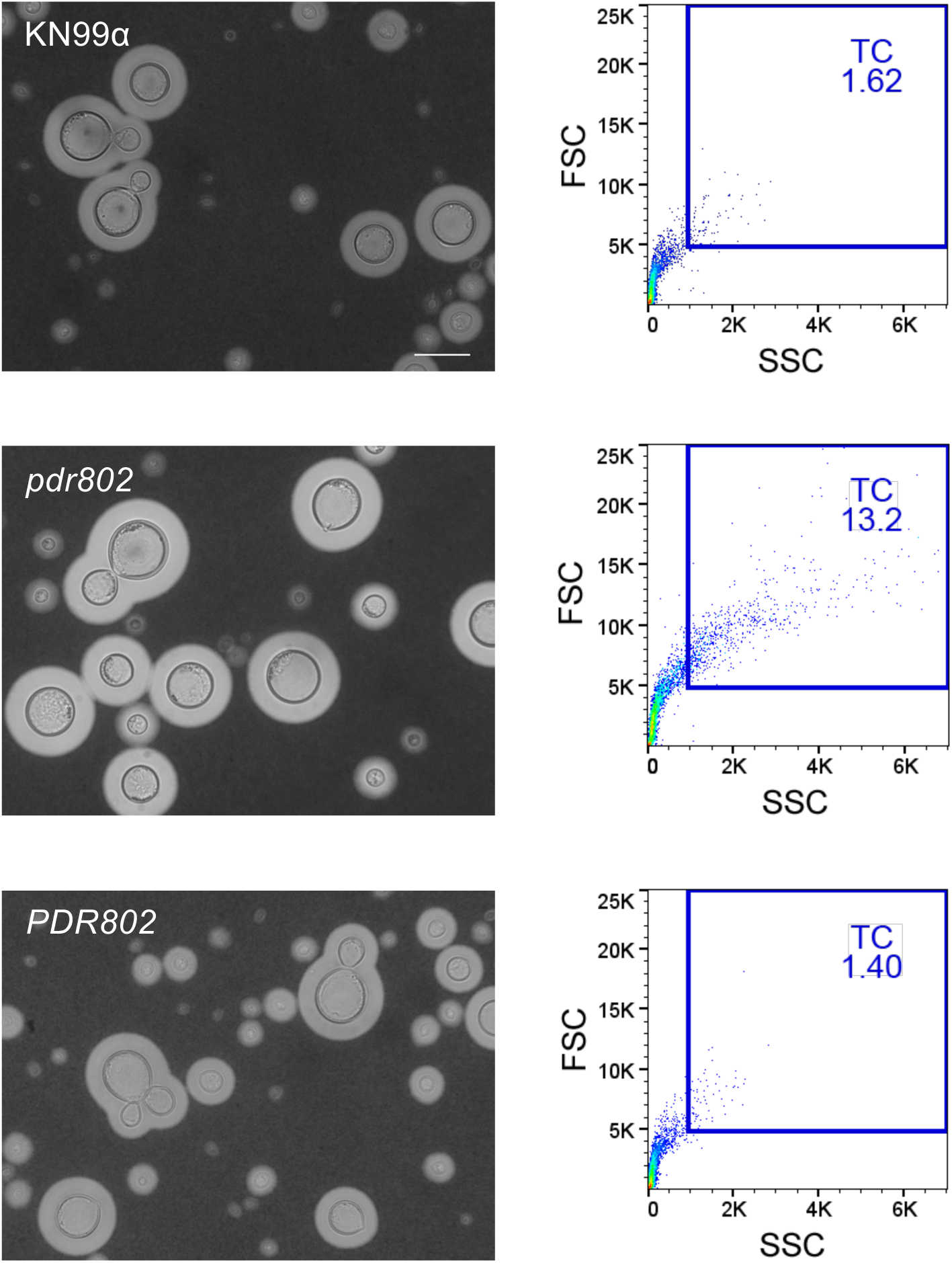
Pdr802 is a negative regulator of Titan cell formation. Left, cultures were subjected to *in vitro* conditions that induce Titan cell formation and imaged with India Ink. All images are to the same scale; scale bar, 10 µm. Images were selected so that each shows multiple examples of Titan cells, not to reflect abundance of this morphotype. Right, the percent of Titan cells (TC) in each culture was quantified using flow cytometry, gated as indicated by the blue square. FSC, forward scatter; SSC, side scatter.

Titan cells are poorly engulfed by host phagocytes (19, 45, 73), which may reflect their increased size as well as alterations in capsule and cell wall (51). We observed this reduced uptake for all strains after growth in conditions that favor Titan cell formation (Figure 7, Titan vs YPD). Also, all strains showed a reduction in phagocytosis after capsule induction in DMEM (Figure 7, DMEM vs YPD), which is not surprising because the capsule is antiphagocytic (31, 73). Notably, the reduction in uptake was greatest for the *pdr802* mutant in both of these conditions, even though it showed normal engulfment when all strains were grown in the control condition (YPD). This is likely because the mutant culture is both hypercapsular and enriched in Titan cells.

**Figure 7.**
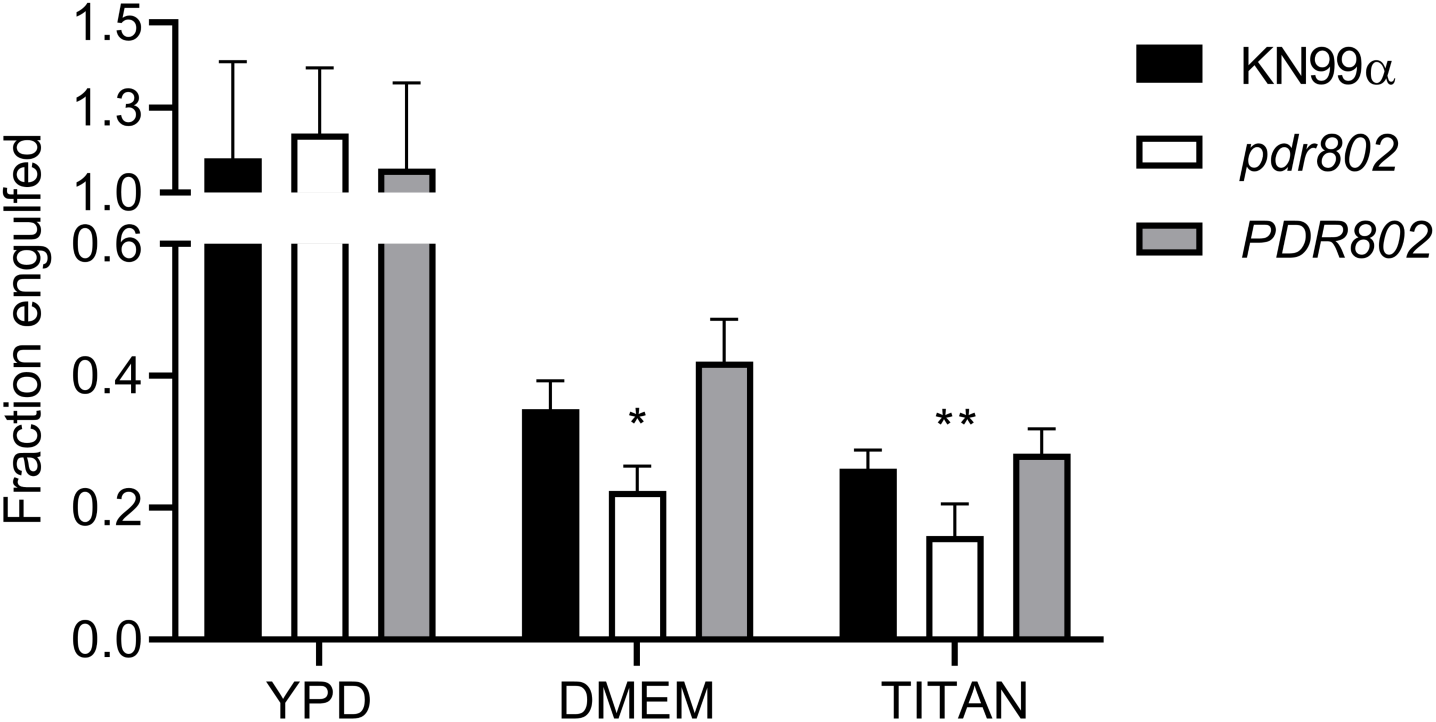
Deletion of *PDR802* affects phagocytosis after growth under conditions that induce capsule and Titan cell formation. The indicated *C. neoformans* strains were grown in YPD (18 h), DMEM (24 h), or Titan-cell induction medium (72 h) and then incubated for 2 h with J774.16 mouse macrophages; host cells were then washed and lysed to assess fungal burden by CFU. Data shown are normalized to the CFU of the initial inoculum. *, p<0.05 and **, p<0.01 compared to KN99α by one-way ANOVA with posthoc Dunnett test.

### Identification of direct, functional targets of Pdr802

To identify direct targets of Pdr802, we performed chromatin immunoprecipitation followed by sequencing (ChIP-Seq). We then compared the DNA sequences immunoprecipitated by anti-mCherry mAb from cells expressing mCherry-Pdr802, which grow similarly to WT (Figure S7A), and untagged cells. Both strains were grown for 24 hours in DMEM at 37°C and 5% CO_2_, as this condition induces *PDR802* expression dramatically compared to standard YPD growth conditions (Figure 2B).

Using 2-fold-enrichment over control as a cutoff value for peaks with adjusted p value <0.05, we identified 656 binding sites for mCherry-Pdr802 in genomic DNA. Of these, 540 occurred within 1,000 bp upstream of transcription start sites (Data Set S1, Sheets 1 and 2), which we used as an approximation of regulatory regions. Application of Discriminative Regular Expression Motif Elicitation (DREME) (74) to this set of upstream regions identified several putative Pdr802 binding motifs, which were highly enriched in GA (TC) (Figure S7B). Notably, the ChIP-seq data also suggested self-regulation of *PDR802*, as has been reported for other cryptococcal TFs (75, 76) (Figure S7C).

To complement our ChIP studies, we determined the set of genes regulated by Pdr802 under host-like conditions by performing RNA-Seq of WT and *pdr802* cells after growth for 24 h in DMEM at 37°C and 5% CO_2_ (Data set 1, Sheet 3). We then used dual-threshold optimization (DTO) to analyze the RNA-seq and ChIP-seq data sets together. This statistical method allowed us to combine the evidence from binding and expression studies to converge on a set of direct and functional TF targets (77). The Pdr802 target genes yielded by this analysis include key players in multiple processes implicated in cryptococcal virulence, including quorum sensing, Titan cell formation, and stress resistance (Data set 1, Sheets 4 and 5).

### Pdr802 represses Titan cell production through regulation of quorum sensing proteins

The most striking phenotype we observed in cells lacking *PDR802* is the marked increase in Titan cell formation. We therefore examined our DTO target list for genes known to influence this phenotype, such as those involved in quorum sensing. Recent studies have shown that the quorum sensing peptide Qsp1 is a negative regulator of Titan cell formation (47, 48); Titan cell formation increases upon deletion of the gene encoding this peptide (*QSP1*) or proteins that mediate its maturation and import (*PQP1* and *OPT1*, respectively). We found that Pdr802 positively regulates *PQP1* and *OPT1* gene expression (Table 1), consistent with its repression of Titan cell formation.

**Table 1.** Pdr802 targets involved in quorum sensing and Titan cell formation.

| <u>Biological process</u> | <u>CNAG</u> | <u>Gene name</u> | <u>ChIP-seq<sup>1</sup></u> | <u>RNA-seq<sup>2</sup></u> | <u>Description</u> |
| --- | --- | --- | --- | --- | --- |
| Quorum sensing | 00150 | *PQP1* | 1.38 | -0.78 | Peptidase |
|  | 03013 | *OPT1* | 1.25 | -0.55 | OPT small oligopeptide transporter |
| Titan cell formation | 05835 | *LIV3* | 1.52 | -0.84 | Transcription factor |
|  | 05785 | *STB4* | 3.33 | -1.71 | Transcription factor |
|  | 05940 | *ZFC3/CQS2* | 2.23 | 0.68 | Transcription factor |

<sup>1</sup>Fold-change for mCherry-Pdr802 compared to WT
<sup>2</sup>Log<sub>2</sub> fold-change for *pdr802* compared to WT

A study of *C. neoformans* cells exposed to Titan cell inducing conditions *in vitro* reported that 562 genes were upregulated in this condition, while 421 genes were downregulated (48). The overlap of these genes with our DTO set of Pdr802-regulated genes included three TF genes *LIV3*, *STB4*, and *ZFC3* (48) (Data Set S2, Sheets 1 and 2). The first two are repressed during Titan cell induction while *ZFC3* (also known as *CQS2*) is induced. Our analysis showed that Pdr802 positively regulates expression of *LIV3* and *STB4*, while it negatively regulates *ZFC3* (Table 1), in concordance with our phenotypic observations of Titan cell formation. Notably, Liv3 and Zfc3 are responsive to the peptide Qsp1 (75, 76) and are important for *C. neoformans* virulence, while Stb4 influences cryptococcal brain infection (67).

### Pdr802 coordinates cryptococcal response to the host environment

*C. neoformans* deploys a variety of proteins to resist the many challenges it experiences upon host entry, which include oxidative and temperature stress. Multiple genes that are central to these responses were identified as direct, functional targets of Pdr802 by our DTO analysis (Table 2). For example, Pdr802 induces the expression of genes whose products detoxify reactive oxygen species (ROS), such as *CAT1*, *CAT2*, and *SOD1* (78, 79), or participate in resistance to these compounds, such as *FZC34*, *MIG1*, and *CCK1* (52, 80, 81) (Table 2). Both the kinase Cck1 (also known as Yck2) and the TF Fzc34 have been implicated in cryptococcal virulence (80, 82).

**Table 2.** Genes regulated by Pdr802 involved in adaptation to the host environment.

| Biological process | CNAG | Gene name | ChIP-seq<sup>1</sup> | RNA-seq<sup>2</sup> | Description |
| --- | --- | --- | --- | --- | --- |
| Oxidative stress resistance | 04981 | *CAT1* | 1.47 | -1.60 | Catalase 1 |
|  | 05256 | *CAT2* | 1.41 | -0.42 | Catalase 2 |
|  | 01019 | *SOD1* | 2.59 | -0.53 | Superoxide dismutase [Cu-Zn] |
|  | 00896 | *FZC34* | 2.22 | -0.82 | Transcription factor |
|  | 06327 | *MIG1* | 1.72 | -0.78 | DNA-binding protein creA |
|  | 00556 | *CCK1* | 1.78 | -0.48 | Casein kinase I |
| Melanin and cell wall formation | 03202 | *CAC1* | 1.60 | -0.61 | Adenylate cyclase |
|  | 01845 | *PKC1* | 2.00 | -0.48 | AGC/PKC protein kinase |
|  | 07724 | *CUF1* | 1.27 | -0.67 | Metal-binding regulatory protein |
|  | 00740 | *SNF5* | 1.28 | -0.70 | Swi/snf chromatin-remodeling subunit |
|  | 01239 | *CDA3* | 2.63 | 1.08 | Chitin deacetylase 3 |
|  | 01230 | *MP98* | 2.74 | 0.96 | Chitin deacetylase 2 |
|  | 00776 | *MP88* | 1.86 | 1.23 | Immunoreactive mannoprotein |
| Growth at 37°C | 00405 | *KIC1* | 1.88 | -0.97 | Ste/ste20/ysk protein kinase |
|  | 03670 | *IRE1* | 1.66 | -1.14 | IRE protein kinase |
| Urease activity | 07448 | *DUR3* | 2.13 | -3.05 | Urea transporter |
|  | 01209 | *FAB1* | 2.29 | -0.49 | 1-phosphatidylinositol-3-P 5-kinase |
|  | 01938 | *KIN1* | 1.61 | -0.34 | CAMK/CAMKL/KIN1 protein kinase |
|  | 01155 | *GUT1* | 2.96 | 0.62 | Glycerol kinase |
|  | 00791 | *HLH1* | 1.71 | -1.09 | Transcription factor |
| Capsule thickness | 02802 | *ARG2* | 1.56 | -0.59 | Inositol/phosphatidylinositol kinase |
|  | 06809 | *IKS1* | 2.28 | -0.44 | IKS protein kinase |
|  | 01905 | *KSP1* | 2.02 | -0.64 | Serine/threonine protein kinase |
|  | 02877 | *FZC51* | 1.75 | -0.69 | Transcription factor |
|  | 07470 | *PDE2* | 2.42 | -1.80 | High-affinity phosphodiesterase |
|  | 03346 | *BZP4* | 3.52 | 1.30 | Transcription factor |
| Calcineurin signaling | 00156 | *CRZ1* | 1.44 | -0.49 | Transcription factor |
|  | 01744 | *HAD1* | 2.26 | 0.47 | Phosphatase |

<sup>1</sup>Fold-change for mCherry-Pdr802 compared to WT<sup>2</sup>Log<sub>2</sub> fold-change for *pdr802* compared to WT

As noted above, melanin has important anti-oxidant properties that promote cryptococcal survival inside the host (16). Under host-like conditions, Pdr802 regulates genes required for melanization, even though it melanizes normally *in vitro*. These genes include *CAC1*, *PKC1*, *CUF1*, and *SNF5* (Table 2). Cac1 is an adenylyl cyclase responsible for cyclic AMP (cAMP) production in *C. neoformans*, which plays a central role in melanin synthesis as well as proper capsule production, mating and virulence (83). The kinase Pkc1 induces production of the laccase (Lac1) that forms melanin and plays a key role in resistance to oxidative and nitrosative stress (84, 85); the TF Cuf1 regulates *LAC1* expression and is important for cryptococcal virulence (86, 87); and *SNF5* is required for full melanization (88). Melanin occurs in the fungal cell wall, which is another key component in fungal stress resistance. Pdr802 is also a direct, functional regulator of several genes whose products influence cell wall glycan content: two chitin deacetylases (Cda3 and Mp98) and the mannoprotein MP88 (Table 2). Changes in mannose and chitin occur in Titan cell walls (51).

Pdr802 positively regulates the expression of several proteins required for yeast growth at 37°C, including the kinases Kic1 and Ire1 (Table 2). Ire1 is a regulator of the cryptococcal Unfolded Protein Response (UPR) pathway and lack of Ire1 or Kic1 impacts *C. neoformans* virulence (80, 89). Pdr802 also modulates cryptococcal urease activity, which is required for dissemination to the central nervous system (CNS) (11, 12), by regulating the urea transporter Dur3 and other proteins that influence urease activity (e.g. the kinases Fab1, Kin1, and Gut1 and the TF Hlh1) (Table 2). Deletion of *FAB1*, *KIN1*, or *HLH1* impair urease activity in *C. neoformans*, while *GUT1* disruption induces it (52, 80).

Above we documented the role of Pdr802 in capsule synthesis, which is dramatically upregulated in the host environment in general and is further increased in cells lacking this TF. We found that Pdr802 is a positive regulator of multiple genes that have been implicated in reducing cryptococcal capsule thickness. These include the kinases Arg2, Iks1, and Ksp1; the TF Fzc51; and the phosphodiesterase Pde2 (Table 2). Notably, null mutants for those genes are hypercapsular, similar to *pdr802* cells (52, 80, 90). Pdr802 is a negative regulator of the TF Bzp4, which, as mentioned above, positively regulates capsule (Table 2) (52).

### Pdr802 regulates calcineurin target genes

The calcineurin signaling pathway is activated by calcium and governs stress response and virulence in *C. neoformans* (91–93). One major mediator of calcineurin signaling is the transcription factor Crz1 mentioned above, which is highly responsive to temperature and influences cryptococcal virulence (57, 62). Upon intracellular calcium influx calcineurin dephosphorylates Crz1, which then translocates to the nucleus and regulates gene expression (57, 63). We found that Pdr802 binds the *CRZ1* gene promoter and positively regulates its expression (Figure 8A, Table 2, and Data Set S2, Sheet 3). Pdr802 also binds and regulates five other genes whose products are dephosphorylated by calcineurin; these include the phosphatase Had1, which is important for cryptococcal cell wall remodeling and virulence (Figure 8B, Table 2, and Data Set S2, Sheet 3) (63, 94).

**Figure 8.**
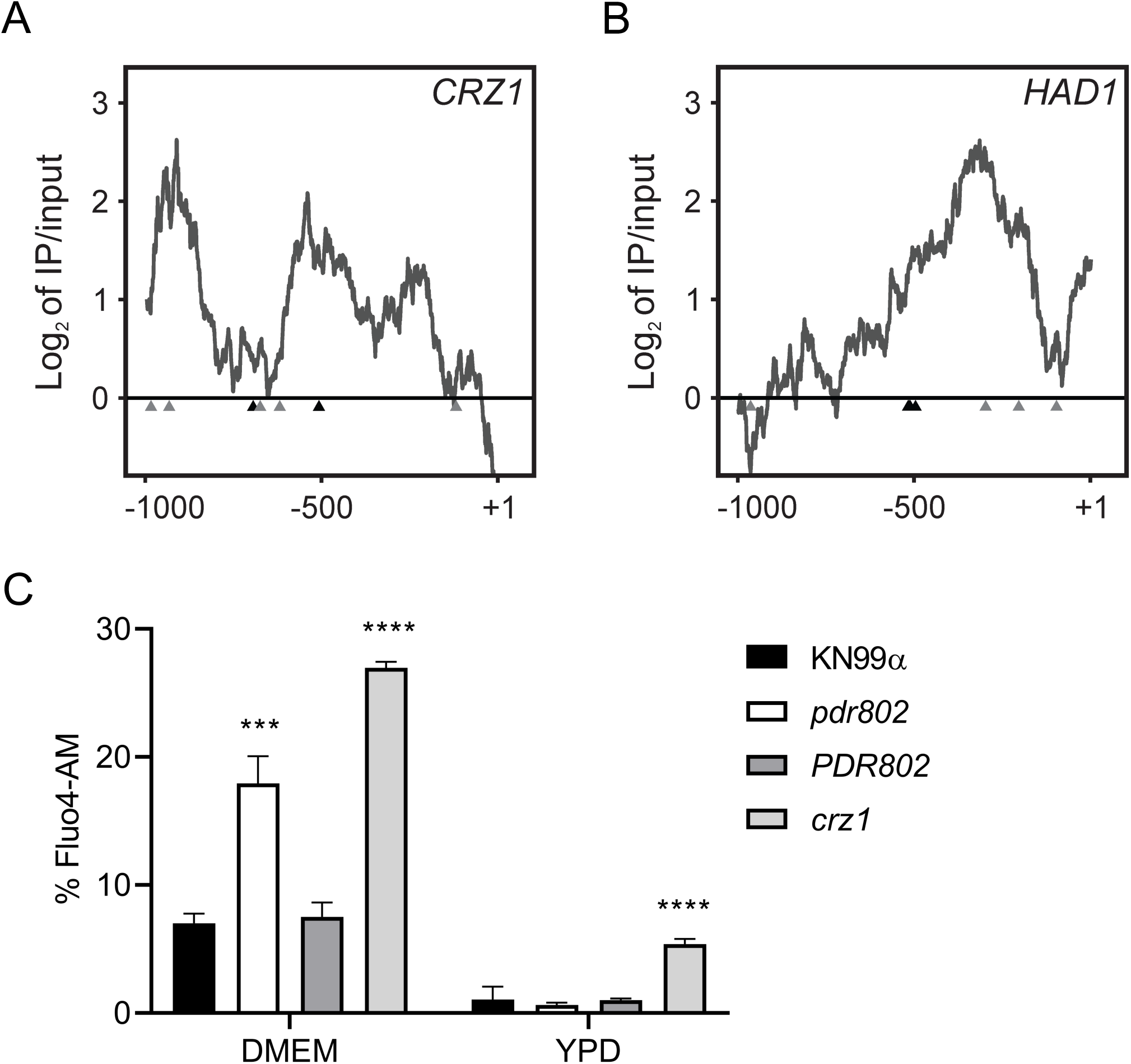
Pdr802 participates in calcineurin signaling. *Panels A-B*. Interactions of Pdr802 with upstream regions of the indicated genes. The ratios (log_2_) of reads from immunoprecipitated (IP) DNA to input DNA were calculated for 1,000 bp upstream of the first coding nucleotide (+1); shown is the difference in these values between tagged and untagged strains. Black triangles, complete Pdr802 DNA-binding motifs (Figure S7B); gray triangles, partial motifs. C. Intracellular calcium measurement by flow cytometry using Fluo-4AM. Each column shows the mean and standard deviation of three biological replicates. ***, p<0.001 and ****, p<0.0001 by one-way ANOVA with posthoc Dunnett test.

Because Crz1 helps maintain normal cryptococcal Ca^2+^ concentrations through the regulation of calcium transporters (57), we wondered about the intracellular calcium levels in *pdr802* cells. We found that after 24 hours of growth in DMEM, the level of cytosolic calcium in the mutant significantly exceeded that of WT or complemented strains (Figure 8C). It was still, however, below that of a *crz1* null mutant, supporting that Pdr802 is not the sole regulator of *CRZ1* expression. Notably, *PDR802* deletion had no effect in rich medium (YPD), which reinforces our hypothesis that Pdr802 acts primarily in host-like conditions. To further explore the relationship of Pdr802 and calcineurin, we compared published gene expression profiles of a calcineurin mutant (57) to our DTO data set. Of the 393 genes that are differently expressed in the calcineurin mutant under thermal stress, 26 are regulated by Pdr802 (Data Set S2, Sheet 4).

## DISCUSSION

We have shown that Pdr802 is a potent regulator of cryptococcal responses to the host environment. In this context, it influences the formation of capsule and Titan cells as well as cellular responses to temperature and oxidative stress, acts as a downstream effector of calcineurin, and modulates calcium availability. The last function is likely achieved through its positive regulation of the transcription factor Crz1, which in turn modulates the calcium transporters Pmc1 and Vcx1 (57). Since calcium ion is a major second messenger in eukaryotic cells, its accumulation in *pdr802* cells affects multiple processes central to host interactions, including stress responses, cell wall integrity, and capsule size (61, 62, 92, 95, 96).

*C. neoformans* dissemination to the brain is the main driver of patient mortality (2). We found that dissemination of *pdr802* cells is significantly impaired, although they do occasionally reach the brain. These observations can be explained by a combination of factors. First, the limited accumulation of the *pdr802* mutant in the lungs, due to factors summarized above, may directly affect dissemination (97). Second, this strain survives poorly in mouse serum, as demonstrated directly by our culture experiments and indirectly by our inability to detect it in the blood of infected mice, even 75 days after infection. The latter might be because the cells do not reach the blood or because they are rapidly eliminated, consistent with previous observations (98). Third, the thick capsules of the *pdr802* mutant reduce its ability to reach the brain. This is true whether fungal entry occurs directly, by the movement of free fungi across the BBB, or indirectly, via a Trojan horse mechanism that requires macrophage uptake (99); such uptake is impeded by enlarged capsules, independent of cell size (31). Fourth, calcium imbalance directly affects cryptococcal transmigration (100). Finally, *pdr802* cells show reduced expression of genes required for urease activity, which promotes *C. neoformans* dissemination to the CNS (11, 12, 100). Interestingly, despite all of these obstacles to dissemination, mutant cells that do reach the brain are able to proliferate to wild-type levels.

Titan cells are a robust and persistent morphotype of *C. neoformans* that contributes to yeast virulence (45). We showed that cells lacking Pdr802 demonstrate increased formation of Titan cells *in vivo* and *in vitro*, suggesting that this TF is a novel repressor of this process. Although Titan cells enhance aspects of cryptococcal pathogenesis (19, 101), their overproduction negatively impacts dissemination to the brain due to their resistance to phagocytosis by macrophages (19, 45) and decreased penetration of biological barriers (19).

Our combined analysis of DNA binding and gene expression data allows us to understand the increase in Titan cell formation that occurs upon deletion of *PDR802*. Under host-like conditions, Pdr802 positively regulates Pqp1, Opt1 and Liv3, all key proteins in the cryptococcal quorum sensing pathway, which represses Titan cell formation (47, 48). In the absence of this TF, quorum sensing is impaired, increasing Titan cell formation. Pdr802 may also indirectly modulate Titan cell formation by regulating other TFs that impact this process, such as Zfc3 (Cqs2) and Stb4.

We know that capsule, a key virulence factor, is typically highly induced in the host or host-like conditions (102). Our studies *in vitro*, *ex vivo*, and *in vivo* show that Pdr802 normally reins in this process. This likely occurs via a combination of Pdr802’s repression of the TF Bzp4, which positively regulates capsule size, and the induction of other factors (e.g. the TF Fzc1, the phosphodiesterase Pde2, and the kinases Ksp1, Arg2, and Iks1) that negatively regulate capsule size (52, 80, 90).

Overall, we found that Pdr802 influences key cryptococcal phenotypes that influence virulence, including quorum sensing, stress responses, Titan cell formation, and capsule production (Figure 9). We have further identified multiple genes that are central in these processes and are directly regulated by Pdr802. Some of these targets are also regulated by calcineurin (e.g. Had1 and Crz1) or by another important TF, Gat201 (e.g. Opt1, Liv3, Zfc3) (60, 75, 76). Finally, the expression of *PDR802* itself is regulated by the TFs Gat201 and Hob1 (67, 76). The crosstalk between all of these regulatory mechanisms remains to be dissected. Nonetheless, it is evident that Pdr802 is critical for both survival in the lung and dissemination to the brain, thus explaining its role in cryptococcal virulence.

**Figure 9.**
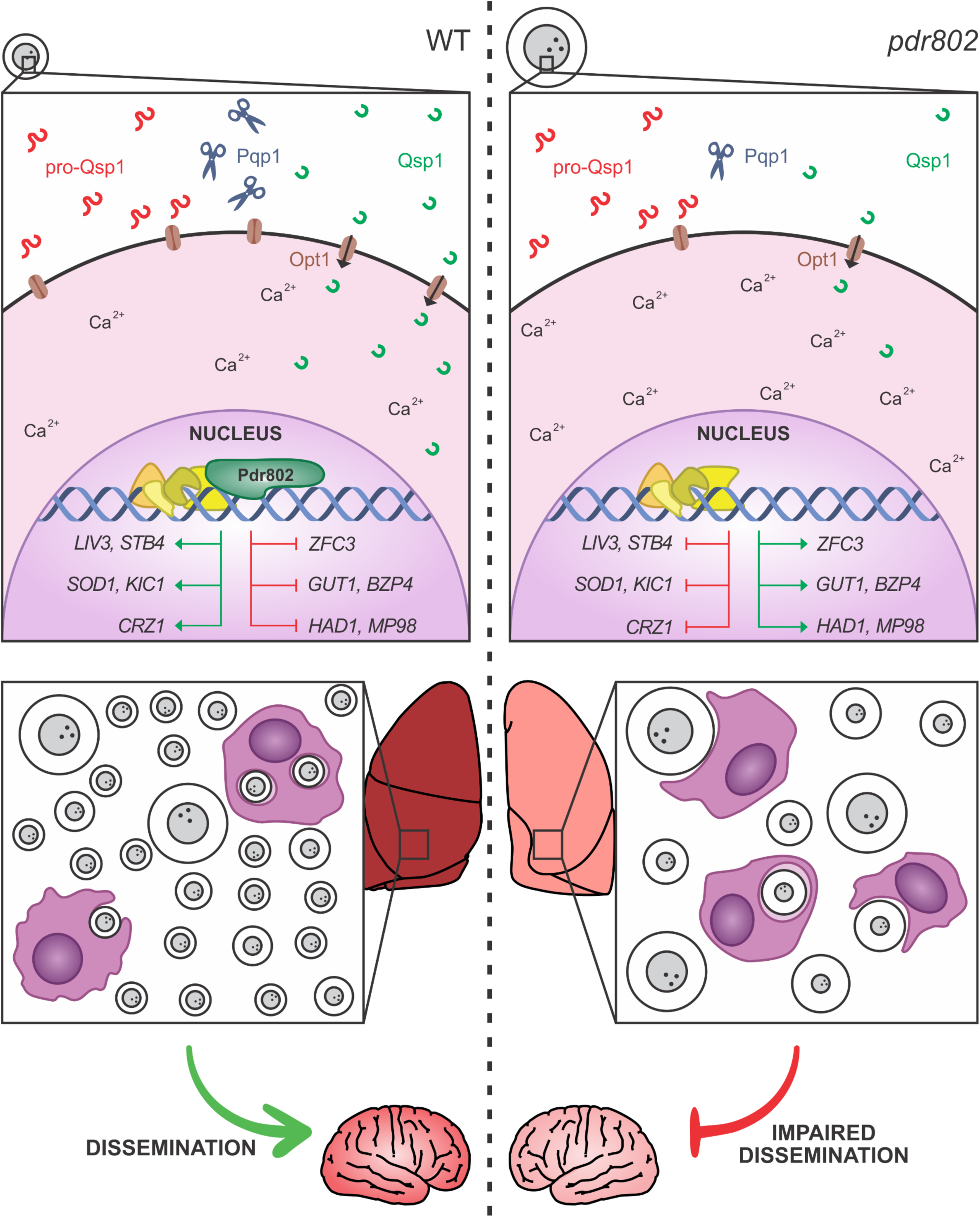
Pdr802 mode of action. *Left panel*. When wild-type *C. neoformans* enters a host, *PDR802* expression is induced and Pdr802 positively regulates elements of the quorum sensing pathway (described in the text) as well as expression of TFs implicated in this pathway (*LIV3*), brain infectivity (*STB4*), and Titan cell production (*ZFC3*). At the same time, Pdr802 regulates two calcineurin targets (*CRZ1* and *HAD1*) and a variety of other genes (see text). Shown are examples of genes involved in the response to oxidative stress (*SOD1*), growth at 37°C (*KIC1*), urease activity (*GUT1*), capsule production (*BZP4*), and cell wall remodeling (*MP98*). *Right panel*. In the absence of these regulatory changes, *pdr802* cells are poorly equipped to survive the stress of the host environment and are subject to increased intracellular calcium levels, dysregulation of capsule production, and impaired stress resistance. As a result, the cryptococcal population in the lung is smaller and is enriched in Titan cells and hypercapsular cells of normal size, both of which demonstrate reduced phagocytosis by host cells and impaired ability to cross biological barriers; these defects reduce dissemination to the central nervous system.

## MATERIALS AND METHODS

### Strain construction and cell growth

We previously reported the *PDR802* deletion mutant (*pdr802*) in the KN99α strain background (103) that was used in this work (39). Complementation of this mutant with the wild-type gene at the native locus (*PDR802*) and construction of a strain that expresses Pdr802 with N-terminal mCherry (mCherry-Pdr802) are detailed in the Supplementary Methods. For all studies, *C. neoformans* strains were inoculated from single colonies into YPD medium (2% [wt/vol] dextrose, 2% [wt/vol] Bacto peptone and 1% [wt/vol] yeast extract in double-distilled water [ddH_2_O]) and grown overnight at 30°C with shaking at 230 rpm before further handling as detailed below. To assess viability during growth in tissue culture medium, overnight cultures were washed with phosphate-buffered saline (PBS), diluted to 10^6^ cells/ml in DMEM (Sigma, D6429), plated (1 ml/well) in triplicate in 24-well plates, and incubated at 37°C and 5% CO_2_. At the indicated times cells were mixed thoroughly, diluted in PBS, and plated on YPD agar (YPD medium, 2% agar [wt/vol]) for assessment of colony-forming units (CFU). To assess viability during growth in mouse serum (prepared as below), YPD-grown cryptococcal cells (10^3^) were incubated in 100 µl of serum in 96-well plates for 24 h at 37°C and 5% CO_2_ and CFU assessed as above.

### Animal experiments

All animal protocols were approved by the Washington University Institutional Animal Care and Use Committee (reference 20170131) or Comissão de Ética no Uso de Animais – CEUA (reference 30936), and care was taken to minimize handling and discomfort. For survival studies, groups of five 4- to 6-week-old female C57BL/6 mice (The Jackson Laboratory) were anesthetized by subcutaneous injection of 1.20 mg ketamine and 0.24 mg xylazine in 120 µl sterile water and intranasally infected with 5 x 10^4^ cryptococcal cells. The mice were monitored and humanely sacrificed when their weight decreased to below 80% of initial weight or if they showed signs of disease, at which point organ burden was assessed. The lungs and brains were harvested, homogenized, diluted and plated on YPD agar. The resulting CFU were enumerated and survival differences were assessed by Kaplan-Meier analysis.

For timed organ burden studies, *C. neoformans* overnight cultures were centrifuged (1,000 x *g* for 3 min), washed with sterile PBS, and resuspended in PBS to 1 x 10^6^ cells/ml. Groups of three 4- to 6-week-old female C57BL/6 mice (Centro Multidisciplinar para Investigação Biológica na Área da Ciência em Animais de Laboratório, CEMIB) were anesthetized as above and intranasally infected with 5 x 10^4^ cryptococcal cells, and monitored as above. At set time points post-infection (see text), mice were sacrificed and fungal burden was assessed from organs (as above) or blood (obtained by cardiac puncture). Organ burden was analyzed by Kruskal-Wallis test with Dunn’s multiple comparison *post hoc* test for each day post-infection.

To assess cryptococcal viability in mouse serum, 6 BALB/c mice were anesthetized with isoflurane and blood was collected from the retro-orbital space using a sterile capillary tube. Collected blood was incubated at 37°C for 30 min and serum was isolated by centrifugation at 1,000 x *g* for 15 min and then heat-inactivated at 56 °C for 30 min.

### Capsule analysis

To qualitatively assess capsule thickness, strains were grown in YPD medium for 16 h, washed with PBS, and 10^6^ cells were incubated in DMEM or mouse serum for 24 h at 37°C and 5% CO_2_. After incubation, cells were fixed in 4% paraformaldehyde, washed three times with PBS, and mixed with similar volumes of India ink for capsule visualization and measurement as previously described (104).

For population-level capsule measurement, *C. neoformans* strains were grown overnight in YPD, washed with PBS, and diluted to 10^6^ cells/ml in DMEM. 150 µl aliquots were then plated in quadruplicate in a poly-L-lysine coated 96-well plate (Fisher 655936) and incubated at 37°C and 5% CO_2_. After 24 hours, the cells were washed with PBS and incubated with 150 µl of a staining mixture (100 µg/ml Calcofluor white to stain cell walls, 50 µg/ml of anticapsular monoclonal antibody 302 conjugated to Alexa Fluor 488 (Molecular probes), and 1.5% goat serum in PBS) for 30 minutes at room temperature in the dark. The cells were washed again with PBS, fixed with 4% formaldehyde for 10 minutes at room temperature, washed with PBS, and each well refilled with 150 µl PBS.

The cells were imaged using a BioTek Cytation 3 imager, which automatically collected 100 images per well in a grid pattern at the well center. Image files were prepared for analysis with the GE InCell Translator and assembled into .xdce image stacks for analysis with the GE INCell Developer Toolbox 1.9. Cell wall and capsule images were first filtered to remove background noise and border objects and then cells were identified using shape-based object segmentation (3-pixel kernel, 50% sensitivity) followed by watershed clump breaking to prevent apparent connectivity caused by incomplete segmentation. Target linking was performed to assign each cell wall object to one capsule object based on known 1:1 pairing and location, generating a target set. Capsule and cell wall object diameters were calculated for each target set (hundreds to thousands per well), and the difference between each pair of measurements was defined as the capsule thickness. Data were normalized by the difference in capsule thickness between uninduced and induced WT cells, which were included in each experiment, and compared to hypercapsular (*pkr1*) (39) and hypocapsular (*ada2*) (105) control strains in each experiment. Capsule sizes were compared by One-Way ANOVA with Dunnett’s multiple comparison *post hoc* test.

To measure capsule thickness of cryptococcal cells grown in the lungs of infected mice, lung homogenates were filtered through a cell strainer with 40 µm pores using a syringe plunger, fixed in 3.7% formaldehyde, and used for India ink staining and measurement as above. For the visualization of KN99α and *PDR802* cells from mouse lungs after 18 days of infection, the tissue was treated with 50 µg/ml DNAse I for 30 min at 37°C.

GXM immunoblotting was conducted as previously described (71). Briefly, 10^6^ cells/ml were grown in DMEM for 24 and 48h. Culture supernatant fractions were then resolved by gel electrophoresis on 0.6% agarose, transferred onto nylon membranes, and probed with 1 μg/ml anti-GXM antibody 302.

### Phenotypic assays

For stress plates, cryptococcal cells were grown overnight in YPD, washed with PBS, and diluted to 10^7^ cells/ml in PBS. Aliquots (3 µl) of 10-fold serial dilutions were spotted on YPD or YNB agar supplemented with various stressors (sorbitol, NaCl, CaCl_2_, LiCl, Congo Red, Calcofluor white, caffeine, SDS, NaNO_2_, H_2_O_2_ and ethanol) in the concentrations indicated in the figures. Melanization was tested on plates made by mixing 10 ml of 2X minimal medium (2 g/L L-asparagine, 1 g/L MgSO_4_ ⋅ 7H_2_O, 6 g/L KH_2_PO_4_, 2 g/L thiamine, 2 mM L-3,4-dihydroxyphenylalanine [L-DOPA] and 0.1% dextrose was added for melanization induction or 0.5% for melanization inhibition) with 10 ml of 2% agar-water per plate. A control strain lacking the ability to melanize was used as a control (*lac1*) (88). For the solid urease assay, 10 µl of a 10^7^ cells/ml suspension in water was plated on Christensen’s urea solid media (1 g/L peptone, 1 g/L dextrose, 5 g/L NaCl, 0.8 g/L KH_2_PO_4_, 1.2 g/L Na_2_HPO_4_, 0.012 g/L phenol red and 15 g/L agar, pH 6.8). Plates were incubated at 30°C or 37°C.

### Titan cells

Titan cell induction was performed in 1x PBS supplemented with 10% heat inactivated Fetal Calf Serum (FCS) for 72 hours at 37°C and 5% CO_2_ as recently described (49) and quantified by flow cytometry as previously reported (47, 48).

### Phagocytosis

J774.16 cells were prepared for uptake experiments by seeding (10^5^ cells/well) in a 96-well plate and incubating in DMEM supplemented with 10% Fetal Bovine Serum (FBS) at 37°C and 5% CO_2_ for 24 h. *C. neoformans* cells were prepared for uptake experiments by inoculating an overnight culture in YPD into either DMEM or Titan cell induction medium (49) and growing at 37°C and 5% CO_2_ for 24 or 72 h, respectively. To initiate the study, cryptococcal cells were washed with PBS and opsonized with anti-capsular antibody 18B7 (1 µg/ml) for 1 h at 37°C while macrophages were activated with 50 nM phorbol myristate acetate (PMA) for 1 h at 37°C and 5% CO_2_; 10^6^ cryptococcal cells were then incubated with the macrophages for 2 h at 37°C and 5% CO_2_. The wells were then washed three times with warm PBS and the macrophages lysed with 0.1% Triton in PBS and plated for CFU as above. Fold-change in CFU was assessed by comparison to the CFU of opsonized cells. One-Way ANOVA with Dunnett’s multiple comparison *post hoc* test was used to compare phagocytosis of *pdr802* and *PDR802* strains with that of KN99α.

### Chromatin Immunoprecipitation (ChIP)

ChIP studies were performed as previously described (39, 105). Briefly, wild type and N-terminal-mCherry-Pdr802 strains were cultivated in DMEM for 24 hours at 37°C and 5% CO_2_. The cells were then fixed with formaldehyde, lysed by mechanical bead-beating, and the cell debris removed by centrifugation. The supernatant fraction was sheared by sonication, centrifuged, and an aliquot was reserved as ‘Input’. The remaining material was incubated with rabbit IgG anti-mCherry antibody (Abcam, ab213511) tethered to protein A sepharose (‘IP’) or sepharose alone (‘Mock’) overnight at 4°C. The beads were then washed, incubated at 65°C to reverse DNA-DNA and DNA-protein crosslinks and the DNA recovered by phenol/chloroform/isoamyl alcohol (25:24:1) extraction, ethanol precipitation, and resuspension in nuclease-free water.

Samples were submitted to the Washington University Genome Technology Access Center for library preparation and DNA samples were sequenced using the Illumina Nextseq platform. The first replicate was sequenced using paired-end 2×75-bp reads and replicates 2 and 3 were sequenced using single-end 75-bp reads; the minimum coverage obtained was ∼16x. The quality of the reads was evaluated by FastQC (106). Fastq files were aligned to the KN99 genome (107) using NextGenMap 0.5.3 (108). SAM files were converted to bam, reads were sorted and indexed, and read duplicates were removed from the final bam files using samtools (109). Samtools was also used to filter out reads with a mapping quality lesser than 20 phreds to guarantee single alignment of the reads. Peaks were called using MACS2 (2.1.1.20160309) (110), filtered by size (maximum threshold 5 kb and no minimum), and annotated using Homer 4.8 (111). The significant peaks were chosen using the cutoff of fold enrichment above 2 and adjusted p value < 0.05 and read coverage of each peak was obtained using Samtools (109). Pdr802 binding motifs were identified using DREME (74); partial motifs were defined as at least 5 consecutive bp of the motif.

### RNA-Seq and Dual-Threshold Optimization (DTO)

RNA from wild-type and *pdr802* cells grown for 24 hours in DMEM (37°C, 5% CO_2_) was isolated and sequenced as previously described (39). Briefly, cDNA samples were sequenced using the Illumina Nextseq platform for single-end 1 x 75 bp reads and read quality was evaluated by FastQC (106). Fastq files were aligned to the KN99 genome (107) using Novoalign (112), SAM files were converted to bam, reads were sorted and indexed, and read duplicates were removed from the final bam files using Samtools (109). The number of reads mapped per gene was calculated using HTSeq (113) and differential gene expression was analyzed with DESeq2 (114), using the Independent Hypothesis Weighting (IHW) package to calculate the adjusted p-values (115). Dual-Threshold Optimization (DTO) analysis was performed as recently described (77). This is a method for simultaneously finding the best thresholds for significance in a TF binding location dataset (e.g. ChIP) and a TF perturbation-response dataset (e.g. RNA-Seq of a TF mutant). It works by trying out all pairs of thresholds for the two datasets, picking the pair that minimizes the probability of the overlap between the bound and responsive gene sets occurring by chance under a null model, and testing the significance of the overlap by comparison to randomly permuted data. Our application of DTO to our ChIP and RNA-Seq data yielded 1455 bound genes, 5186 responsive genes, and 1167 genes that were both bound and responsive. Based on DTO, Pdr802 has an acceptable convergence from binding and perturbation, with p value < 0.01 from the random permutation test and minimum expected FDR less than or equal to 20% at 80% sensitivity. In addition to requiring a statistically significant overlap between the ChIP-Seq and RNA-Seq gene sets, we filtered out any genes for which traditional differential expression analysis yielded an adjusted p-value ≤ 0.15 or absolute log_2_ of fold change ≥ 0.3, leaving 380 bound targets.

### Intracellular calcium measurement

To measure intracellular free Ca^2+^, yeast cells were cultured overnight in YPD at 30°C with shaking, washed three times with deionized water, diluted to 10^6^ cells/ml in DMEM (Sigma, D6429), plated (1 ml/well) in triplicate in 24-well plates, and incubated at 37°C and 5% CO_2_ for 24 hours. At the indicated times, cells were mixed thoroughly, diluted in PBS containing 2 μM Fluo4-AM (Thermo Fisher), incubated at 30°C for 30 min, and analyzed using flow cytometry. The overnight culture was used as a control and treated as above.

## Data availability

ChIP-seq and RNA-seq data files are available at the NCBI Gene Expression Omnibus under accession numbers GSE153134 and GSE162851, respectively.

## ACKNOWLEDGMENTS

We appreciate helpful discussions with Charley Christian Staats, Augusto Schrank, and members of the Doering, Kmetzsch and Brent labs. We are grateful for assistance from Thomas Hurtaux, Eamim Squizani, and Julia Sperotto with mouse experiments; Guohua Chen with spotting assays; Jessica Plaggenberg with library preparation and sequencing; Chase Mateusiak with RNA-seq data analysis; and Sandeep Acharya for performing the DTO analysis. We thank Marilene Henning Vainstein and her lab for providing support with *in vivo* experiments. We also thank Liza Miller for comments on the manuscript and Arturo Casadevall for providing the antibody anti-GXM (18B7).

These studies were supported by National Institutes of Health grant AI087794 to TLD and MRB; National Institutes of Health grants AI36688 and AI140979 to TLD; and grants from Coordenação de Aperfeiçoamento de Pessoal de Nível Superior (CAPES, Brazil), Conselho Nacional de Desenvolvimento Científico e Tecnológico (CNPq, Brazil – Grant number 310510/2018-0), and Fundação de Amparo à Pesquisa do Estado do Rio Grande do Sul (FAPERGS, Brazil) to LK. CAPES fully supported JCVR during her studies in Brazil; her studies in the United States were partially supported by this source (Advanced Network of Computational Biology - RABICÓ - Biocomputational Grant 23038.010041/2013-13).

## AUTHORS CONTRIBUTION

Conceived and designed experiments: J.C.V.R., D.P.A., A.L.C., M.R.B., L.K. and T.L.D.;

Performed experiments: J.C.V.R., D.P.A., H.M., H.B., and A.L.C;

Analyzed data: J.C.V.R., D.P.A., A.L.C., M.R.B., L.K. and T.L.D.;

Contributed reagents and materials: M.R.B., L.K. and T.L.D.;

Drafted the paper: J.C.V.R, L.K. and T.L.D.

Revised the paper: J.C.V.R., D.P.A., M.R.B., L.K. and T.L.D.

## SUPPLEMENTAL FIGURE LEGENDS

**Figure S1. Mutant strain construction and confirmation.** A. Scheme for generating *C. neoformans* strains in the KN99α background (middle) that either lack *PDR802* (*pdr802,* top) or encode a tagged copy of the protein (*mCherry-PDR802*). B. Qualitative analysis of gene expression in Panel A strains and the complemented *pdr802* mutant (*PDR802*). Cryptococcal mRNA isolated from cells grown in DMEM (37°C, 5% CO_2_, 24 hours) was used to generate cDNA; from this, segments of the genes indicated at the left were amplified using the primers listed in Data Set S2, Sheet 5, and the products were analyzed by agarose gel electrophoresis. Fragment sizes (in bp) are indicated at right and the ladder bands shown are 400, 500, 650, 850 and 1000 bp for the top panel; 200, 300, 400, 500, and 650 bp for the middle panel; and 100, 200, and 300 bp for the bottom panel. C. Quantitative analysis of *PDR802* expression. Samples of RNA isolated as in B were analyzed for *PDR802* expression by qRT-PCR. All results were normalized to *ACT1* expression. Each symbol represents a biological replicate, with the mean and standard deviation also shown. ***, p<0.001 compared to KN99α by one-way ANOVA with posthoc Dunnett test.

**Figure S2. Organ burdens.** A. Mean +/− SD values of total colony-forming units (CFU) in the indicated tissue of mice from the Figure 1 survival curve are shown. Each point is the average value for a single animal at the time of death. For *pdr802* infections, red circles represent mice sacrificed at days 65 and 69, while blue circles represent mice sacrificed at the end of the study (day 100). B. Mean +/− SD of total colony-forming units (CFU) in the blood and brain at the indicated times post-infection. C. Mean +/− SD of total colony-forming units (CFU) in the lung, blood and brain 75 days after infection with *pdr802*. Each color represents one mouse.

**Figure S3. Characterization of *pdr802* cells.** *Panels A-C*.10-fold serial dilutions of WT, *pdr802*, and *PDR802* cells were plated on the media shown and incubated at 30°C (A), 37°C (B), or 37°C in the presence of 5% CO_2_ (C). Nitrosative (NaNO_2_) and oxidative (H_2_O_2_) stress plates were prepared with YNB medium and melanization plates containing L-DOPA were prepared as in the Methods; all other plates were prepared with YPD medium. *lac1*, a control strain lacking the ability to melanize (88). D. Urease activity of the indicated strains was evaluated using Christensen’s urea solid medium (see Methods) at the indicated temperatures. *ure1*, a control strain that does not produce urease (17).

**Figure S4. Growth curves and capsule shedding.** *Panels A-B*. Growth of the strains indicated in YPD at 30°C (A) or DMEM at 37°C and 5% CO_2_ (B) was assessed by OD_600nm_ at the times shown. C. Conditioned medium from the indicated strains was probed for the presence of GXM after growth in DMEM for 24 or 48 hours. Equal volumes of culture supernatant were analyzed without normalization to cell density. Immunoblotting was performed using the anti-GXM monoclonal antibody 302.

**Figure S5. Semi-automated assay for cryptococcal capsule imaging.** A. Schematic of applying this method to cryptococcal cells induced to form capsule by growth in DMEM (37°C, 5% CO_2_) for 24 h, followed by cell wall and capsule staining. Thousands of cells may be imaged per well and analyzed automatically with software that annotates and measures the capsule (annotated on the micrograph in blue) and cell wall (annotated in bright green). See Methods for details. B. Capsule size distribution of WT cells after induction. Capsule thickness for each cell is the difference between the paired diameters of the cell wall and capsule, which is plotted here with reference to the mean value. C and D. Mean and SD (C) and cumulative percentage (D) analysis of WT compared to hyper and hypocapsular control strains (here *pkr1* and *ada2*, respectively). Capsule thickness is in arbitrary units, related to the pixels measured. E. The time required to analyze the capsule thickness of 1,000 cells by this method compared to manual assessment of India ink images.

**Figure S6. *PDR802* deletion induces Titan cell formation.** Mean +/− SD of (A) total cell diameter and (B) the ratio of total cell to cell body diameters (diameter ratio), assessed by measuring at least 50 cells per strain with ImageJ. **, p<0.01 and ****, p<0.0001 for comparison of *pdr802* to KN99α or *PDR802* by one-way ANOVA with posthoc Dunnett test for each day post-infection. C. Percent of Titan cells in the indicated strain, evaluated using various published parameters: cell body diameter above 10 or 15 µm (20) or total cell diameter above 30 µm (43).

**Figure S7. Pdr802 strain viability, putative DNA-binding motifs, and self-regulation.** A. The indicated strains were grown in DMEM at 37°C and 5% CO_2_ for the times shown and samples were tested for their ability to form colonies on YPD medium. Plotted is the fold-change in CFU relative to the initial culture. B. Putative Pdr802-binding motifs determined using DREME (74). Primary and secondary hits are shown for analysis of 1,000 bp upstream of the initiating ATG. C. Pdr802 self-regulation. The ratios (log_2_) of reads from immunoprecipitated (IP) DNA to reads from input DNA were calculated for 1,000 bp upstream of the first coding nucleotide (+1) of *PDR802*; shown is the difference in these values between tagged and untagged strains. Red triangles, complete Pdr802 DNA-binding motifs (Figure S7B); blue triangles, partial motifs.

**Data Set S1. Pdr802 ChIP-Seq, RNA-Seq, and Dual-Threshold Optimization (DTO) data.** Sheet 1, peaks that occur in gene promoter regions that showed ≥2-fold enrichment when Chip-Seq was performed on strains expressing tagged versus untagged Pdr802, with annotation. Sheet 2, ChIP-Seq primary data of Pdr802-specific peaks. Sheet 3, RNA-Seq data. Sheet 4, DTO filtered data. Sheet 5, DTO primary data. Sheet 6, ChIP-Seq data of all peaks in mCherry-Pdr802 samples. Sheet 7, ChIP-Seq data of all peaks in untagged (WT) control samples.

**Data Set S2. Pdr802 target analysis and primers used in this study.** Sheet 1, Pdr802 regulated genes that are down-regulated during Titan cell formation *in vitro* (48). Sheet 2, Pdr802 regulated genes that are up-regulated during Titan cell formation *in vitro* (48). Sheet 3, targets regulated by Pdr802 that are also dephosphorylated by calcineurin (63). Sheet 4, the intersection of Pdr802 targets and genes that are Crz1-independent calcineurin targets under conditions thermal stress (55). Sheet 6, primers used in this study. Fold enrichment and adjusted p (q) values throughout are for the DTO results.

**Supplementary Methods.**

