## Supplemental figures and methods for "The transcription factor Pdr802 regulates Titan cell formation, quorum sensing, and pathogenicity of *Cryptococcus neoformans*"

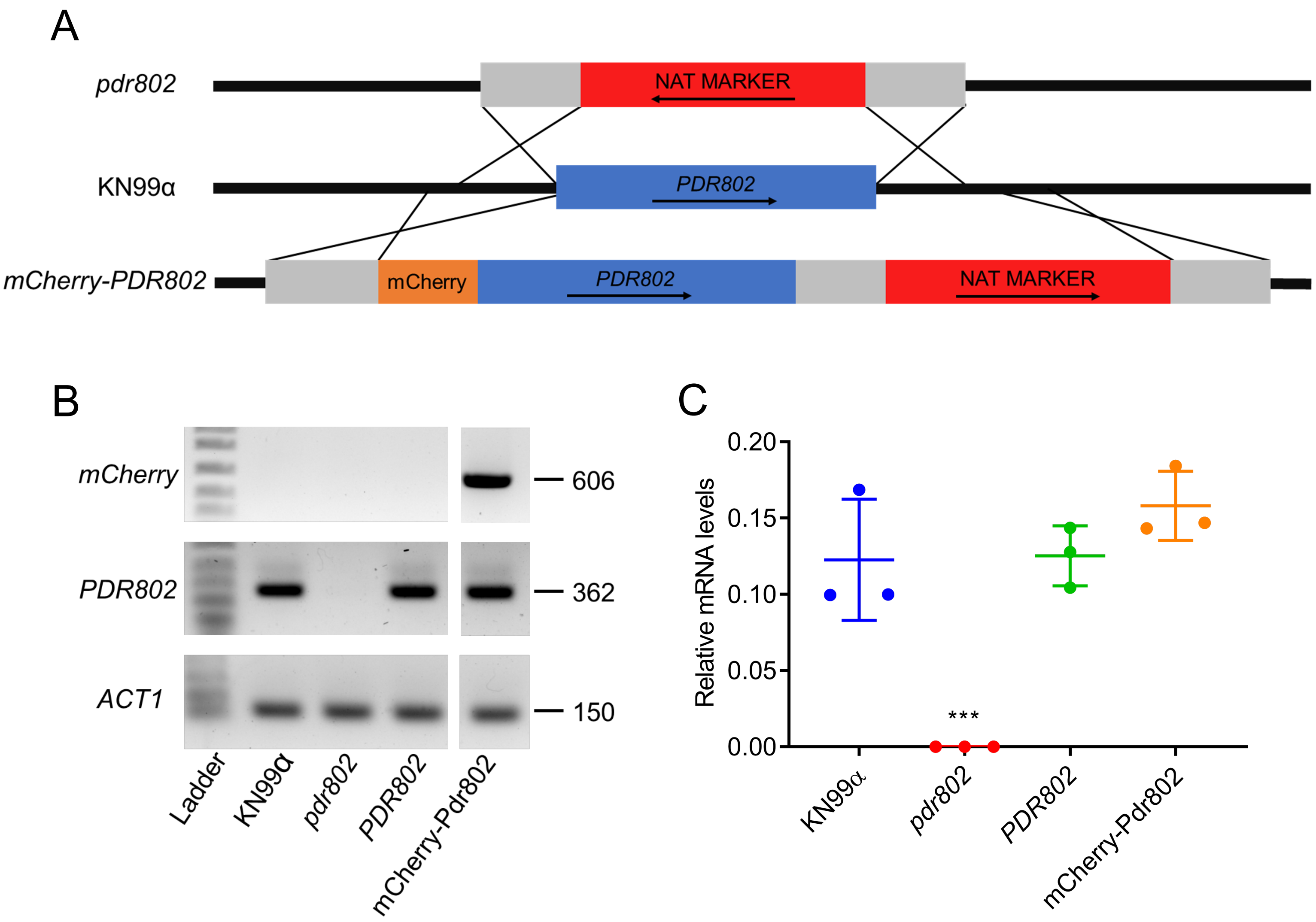


**Figure S1. Mutant strain construction and confirmation.** A. Scheme for generating *C. neoformans* strains in the KN99α background (middle) that either lack *PDR802* (*pdr802,* top) or encode a tagged copy of the protein (*mCherry-PDR802*). B. Qualitative analysis of gene expression in Panel A strains and the complemented *pdr802* mutant (*PDR802*). Cryptococcal mRNA isolated from cells grown in DMEM (37°C, 5% $CO_2$, 24 hours) was used to generate cDNA; from this, segments of the genes indicated at the left were amplified using the primers listed in Data Set S2, Sheet 5, and the products were analyzed by agarose gel electrophoresis. Fragment sizes (in bp) are indicated at right and the ladder bands shown are 400, 500, 650, 850 and 1000 bp for the top panel; 200, 300, 400, 500, and 650 bp for the middle panel; and 100, 200, and 300 bp for the bottom panel. C. Quantitative analysis of *PDR802* expression. Samples of RNA isolated as in B were analyzed for *PDR802* expression by qRT-PCR. All results were normalized to *ACT1* expression. Each symbol represents a biological replicate, with the mean and standard deviation also shown. ***, p<0.001 compared to KN99α by one-way ANOVA with posthoc Dunnett test.

FIGURE S2

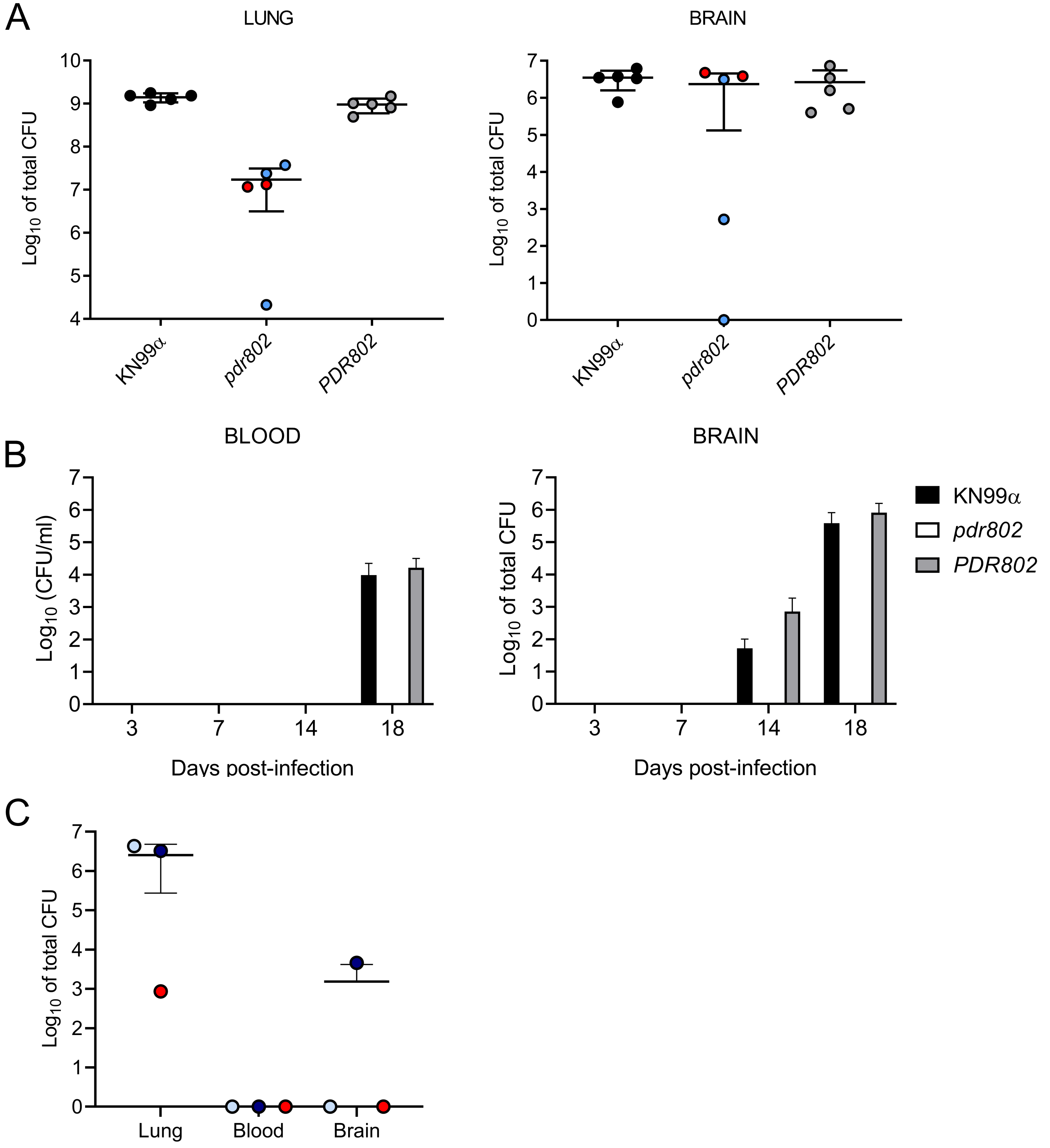


**Figure S2. Organ burdens.** A. Mean +/- SD values of total colony-forming units (CFU) in the indicated tissue of mice from the Figure 1 survival curve are shown. Each point shows the average value for a single animal at the time of death. For *pdr802* infections, red circles represent mice sacrificed at days 65 and 69, while blue circles represent mice sacrificed at the end of the study (day 100). B. Mean +/- SD of total colony-forming units (CFU) in the blood and brain at the indicated times post-infection. C. Mean +/- SD of total colony-forming units (CFU) in the lung, blood and brain 75 days after infection with *pdr802*. Each color represents one mouse.

### FIGURE S3

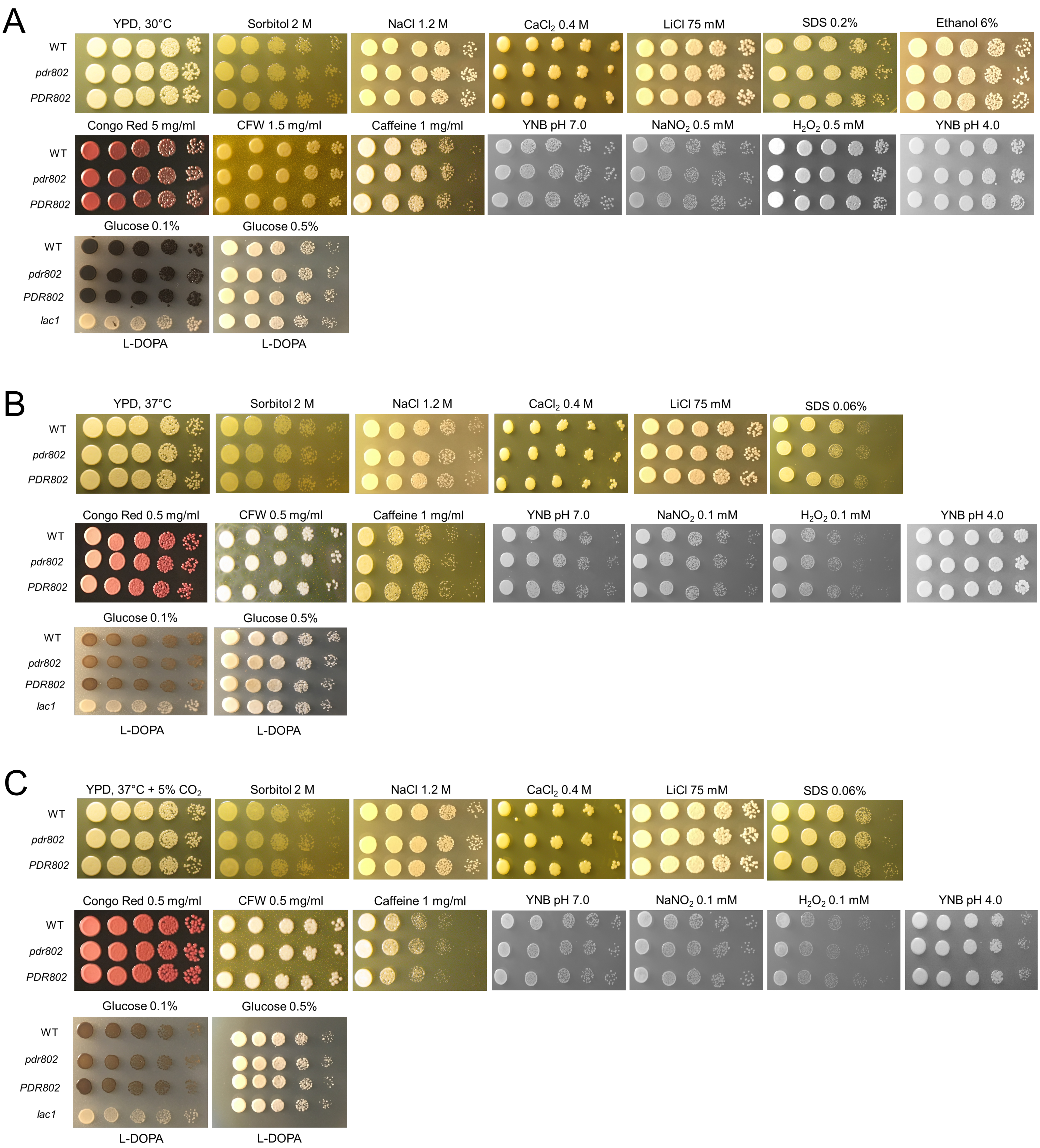
A
YPD, 30°C
Sorbitol 2 M
NaCl 1.2 M
CaCl2 0.4 M
LiCl 75 mM
SDS 0.2%
Ethanol 6%
WT
pdr802
PDR802
Congo Red 5 mg/ml
CFW 1.5 mg/ml
Caffeine 1 mg/ml
YNB pH 7.0
NaNO2 0.5 mM
H2O2 0.5 mM
YNB pH 4.0
Glucose 0.1%
Glucose 0.5%
lac1
L-DOPA
L-DOPA
B
YPD, 37°C
Sorbitol 2 M
NaCl 1.2 M
CaCl2 0.4 M
LiCl 75 mM
SDS 0.06%
Congo Red 0.5 mg/ml
CFW 0.5 mg/ml
Caffeine 1 mg/ml
YNB pH 7.0
NaNO2 0.1 mM
H2O2 0.1 mM
YNB pH 4.0
Glucose 0.1%
Glucose 0.5%
L-DOPA
L-DOPA
C
YPD, 37°C + 5% CO2
Sorbitol 2 M
NaCl 1.2 M
CaCl2 0.4 M
LiCl 75 mM
SDS 0.06%
Congo Red 0.5 mg/ml
CFW 0.5 mg/ml
Caffeine 1 mg/ml
YNB pH 7.0
NaNO2 0.1 mM
H2O2 0.1 mM
YNB pH 4.0
Glucose 0.1%
Glucose 0.5%
L-DOPA
L-DOPA

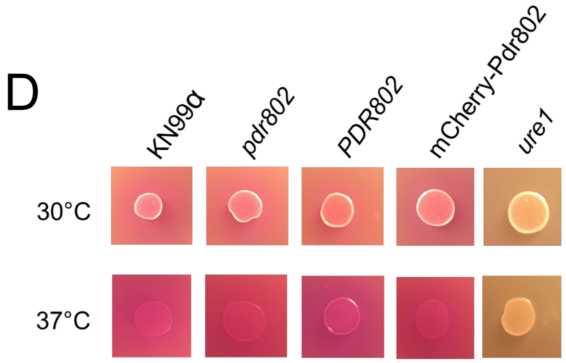


**Figure S3. Characterization of *pdr802* cells.** *Panels A-C*.10-fold serial dilutions of WT, *pdr802*, and *PDR802* cells were plated on the media shown and incubated at 30°C (A), 37°C (B), or 37°C in the presence of 5% $CO_2$ (C). Nitrosative ($NaNO_2$) and oxidative ($H_2O_2$) stress plates were prepared with YNB medium and melanization plates containing L-DOPA were prepared as in the Methods; all other plates were prepared with YPD medium. *lac1*, a control strain lacking the ability to melanize (88). D. Urease activity of the indicated strains was evaluated using Christensen's urea solid medium (see Methods) at the indicated temperatures. *ure1*, a control strain that does not produce urease (17).

FIGURE S4

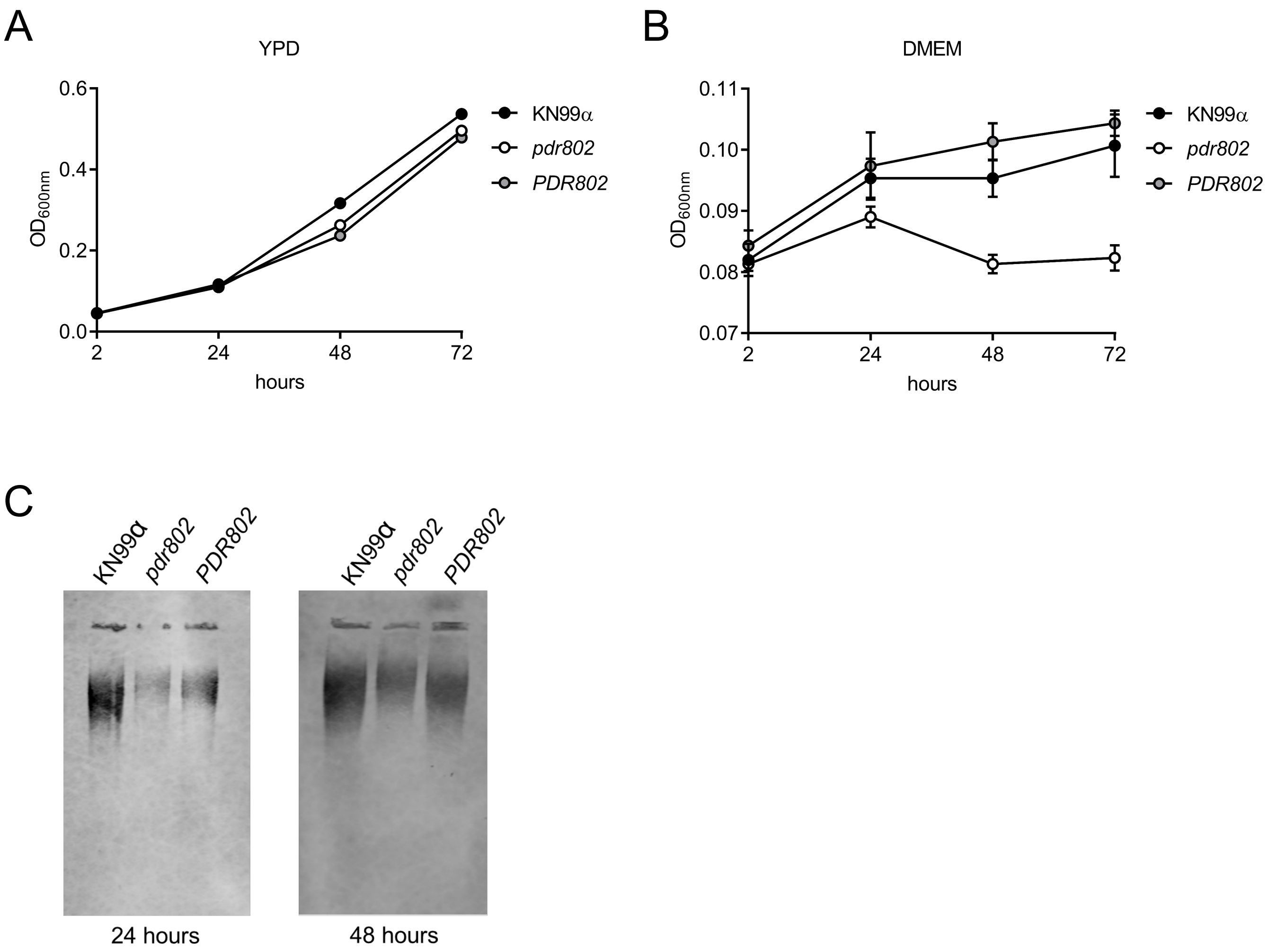


**Figure S4. Growth curves and capsule shedding.** *Panels A-B*. Growth of the indicated strains in YPD at 30°C (A) or DMEM at 37°C and 5% $CO_2$ (B) was assessed by $OD_{600nm}$ at the times shown. C. Conditioned medium from the indicated strains was probed for the presence of GXM after growth in DMEM for 24 or 48 hours. Equal volumes of culture supernatant were analyzed without normalization to cell density. Immunoblotting was performed using the anti-GXM monoclonal antibody 302.

FIGURE S5

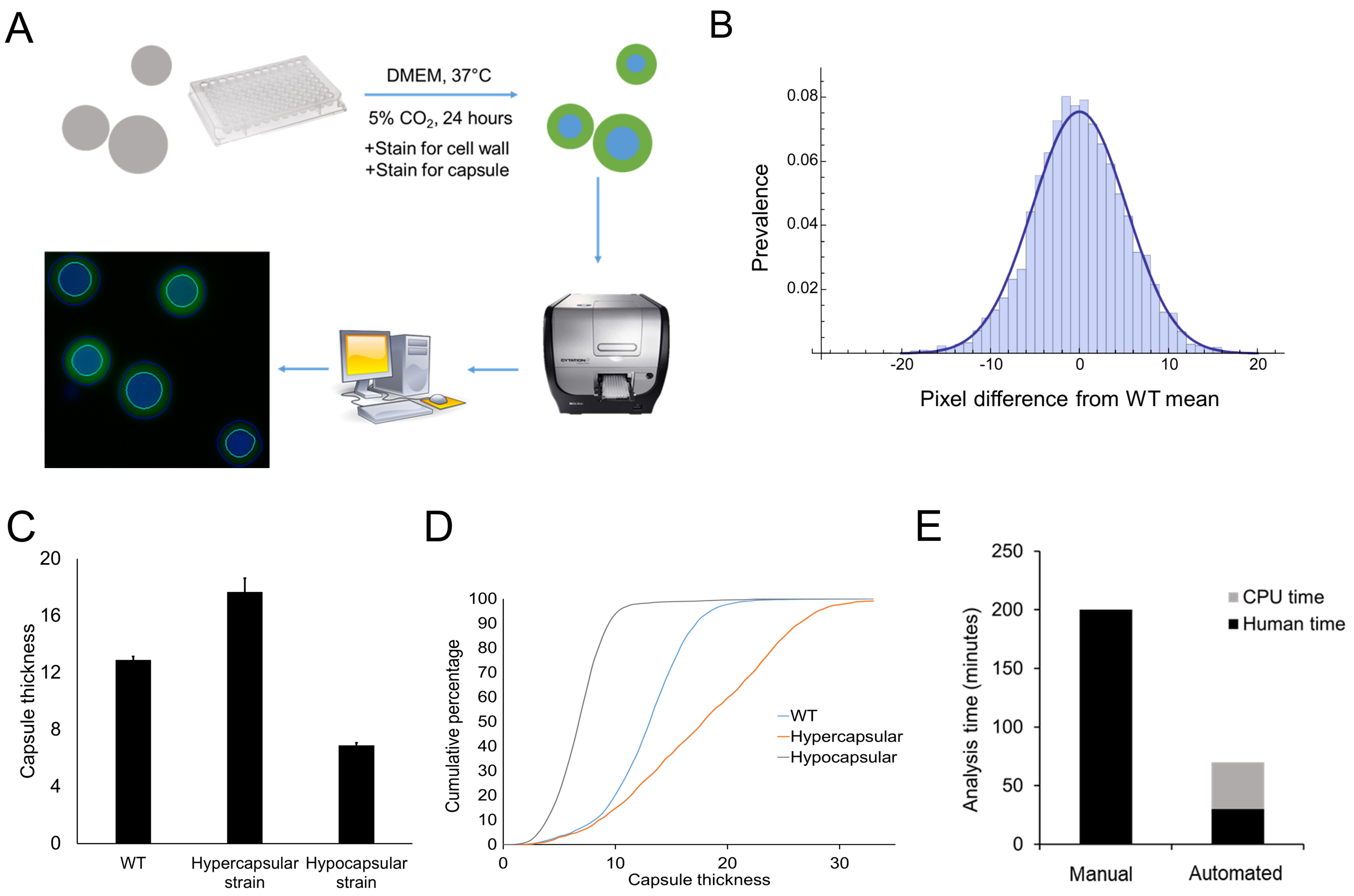


**Figure S5. Semi-automated assay for cryptococcal capsule imaging.** A. Schematic of applying this method to cryptococcal cells induced to form capsule by growth in DMEM (37°C, 5% $CO_2$) for 24 h, followed by cell wall and capsule staining. Thousands of cells may be imaged per well and analyzed automatically with software that annotates and measures the capsule (annotated on the micrograph in blue) and cell wall (annotated on the micrograph in bright green). See Methods for details. B. Capsule size distribution of WT cells after induction. Capsule thickness for each cell is the difference between the paired diameters of the cell wall and capsule, which is plotted here with reference to the mean value. C and D. Mean and SD (C) and cumulative percentage (D) analysis of WT compared to hyper and hypocapsular control strains (here *pkr1* and *ada2*, respectively). Capsule thickness is in arbitrary units, related to the pixels measured. E. The time required to analyze the capsule thickness of 1,000 cells by this method compared to manual assessment of India ink images.

FIGURE S6

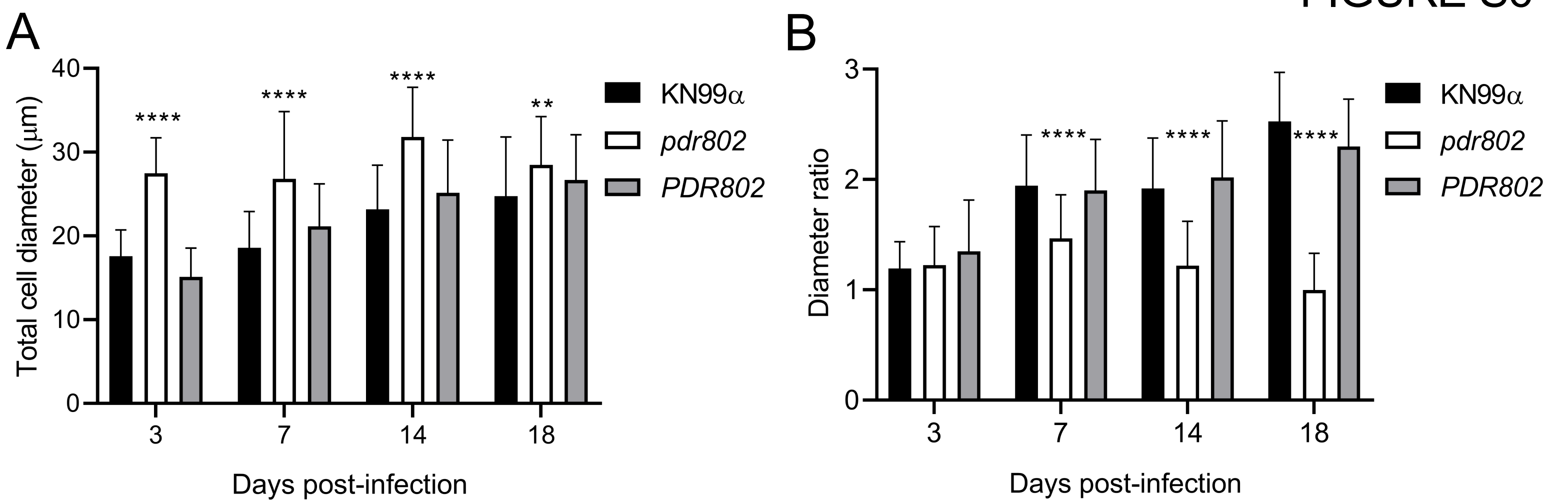


C

| 3 DPI | KN99α | *pdr802* | *PDR802* |
|---|---|---|---|
| Cell body diameter ≥ 10 μm | 20 | 86.7 | 11.1 |
| Cell body diameter ≥ 15 μm | 0 | 22.2 | 0 |
| Total cell diameter ≥ 30 μm | 0 | 31.1 | 0 |
| **7 DPI** | **KN99α** | ***pdr802*** | ***PDR802*** |
| Cell body diameter ≥ 10 μm | 8.5 | 57.4 | 8.5 |
| Cell body diameter ≥ 15 μm | 0 | 19.1 | 2.1 |
| Total cell diameter ≥ 30 μm | 0 | 31.9 | 2.1 |
| **14 DPI** | **KN99α** | ***pdr802*** | ***PDR802*** |
| Cell body diameter ≥ 10 μm | 27.1 | 87.1 | 24.3 |
| Cell body diameter ≥ 15 μm | 2.9 | 42.9 | 4.3 |
| Total cell diameter ≥ 30 μm | 12.9 | 57.1 | 25.7 |
| **18 DPI** | **KN99α** | ***pdr802*** | ***PDR802*** |
| Cell body diameter ≥ 10 μm | 9.1 | 89.4 | 15.1 |
| Cell body diameter ≥ 15 μm | 0 | 53.0 | 1.5 |
| Total cell diameter ≥ 30 μm | 21.2 | 43.9 | 27.3 |

**Figure S6. *PDR802* deletion induces Titan cell formation.** Mean +/- SD of (A) total cell diameter and (B) the ratio of total cell to cell body diameters (diameter ratio), assessed by measuring at least 50 cells per strain with ImageJ. **, $p<0.01$ and ****, $p<0.0001$ for comparison of *pdr802* to KN99α or *PDR802* by one-way ANOVA with posthoc Dunnett test for each day post-infection. C. Percent of Titan cells in the indicated strain, evaluated using various published parameters: cell body diameter above 10 or 15 μm (20) or total cell diameter above 30 μm (43).

### FIGURE S7

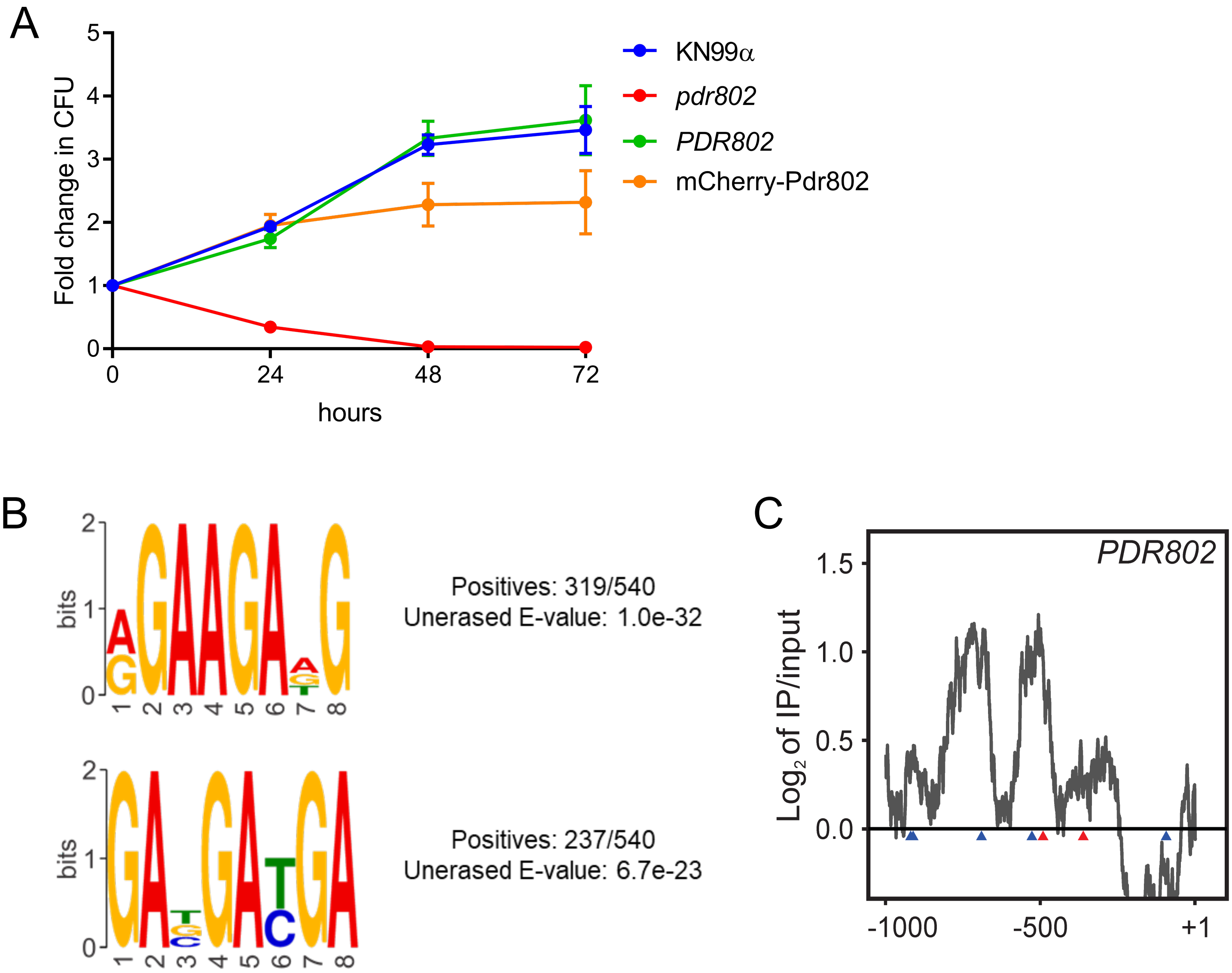


**Figure S7. Pdr802 strain viability, putative DNA-binding motifs, and self-regulation.** A. The indicated strains were grown in DMEM at 37°C and 5% $CO_2$ for the times shown and samples were tested for their ability to form colonies on YPD medium. Plotted is the fold-change in CFU relative to the initial culture. B. Putative Pdr802-binding motifs determined using DREME (76). Primary and secondary hits are shown for analysis of 1,000 bp up-stream of the initiating ATG. C. The ratios ($log_2$) of reads from immunoprecipitated (IP) DNA to reads from input DNA were calculated for 1,000 bp upstream of the first coding nucleotide (+1) of *PDR802*; shown is the difference in these values between tagged and untagged strains. Red triangles, complete Pdr802 DNA-binding motifs (Figure S7B); blue triangles, partial motifs.

SUPPLEMENTARY METHODS

*Strain construction*

To complement the *pdr802* deletion mutant at its native locus, we used a split marker strategy (1) to replace the nourseothricin marker with the original *PDR802* sequence (including its native promoter and terminator) followed by a geneticin (G418) resistance marker. To do this we PCR amplified the *PDR802* sequence (4131 bp) with primers JR01/02 and the G418 resistance marker (1627 bp) with primers JR03/04 (see the Data Set S2, Sheet 5 for primer sequences); both amplicons were then gel purified and cloned in tandem into NdeI-digested pUC19, using Gibson Assembly Master Mix (2), to form pUC19+PDR802+G418. This plasmid was digested with NdeI and BssHII to release a 5492-bp fragment containing *PDR802* and a 5' portion of the G418 marker. The 3' portion of the G418 resistance marker (750 bp) was amplified using primers JR05/06 and fused by PCR to a region 3' of *PDR802* (1017 bp; amplified with JR07/08) to yield a 1767 bp product. These fragments were used in biolistic transformation of *pdr802* cells as in ref (3).

To tag Pdr802 with mCherry at the N-terminus, we first fused approximately 1.2 kb upstream of the start codon of *PDR802,* a sequence encoding mCherry, the *PDR802* coding sequence with 500 bp of downstream sequence, and a NAT resistance marker sequence. To generate these segments the *PDR802* promoter (1286 bp) was amplified with primers JR09/10; the mCherry sequence (714 bp) with JR11/12; the *PDR802* coding sequence (2957 bp) with JR13/14; and the NAT marker cassette (1721 bp) with primers JR15/16. All fragments were gel purified and cloned as above into pUC19 digested with Sac1 and Nde1, to form pUC19+mCherry+Pdr+NAT. This was digested with NdeI and BssHII to release a 5559 bp fragment. The 3' portion of the NAT resistance marker (832 bp) was amplified with primers JR17/18 and fused by PCR to a region 3' of *PDR802* amplified with JR19/20 (1064 bp) to form a 1896-bp product. These fragments were used in biolistic transformation of KN99α cells as above. All strains were confirmed by PCR and sequencing, and further analyzed by RT-PCR and qRT-PCR (see Supplemental Figure 1).

*qRT-PCR*

RNA from cryptococcal cells grown in DMEM at 37°C with 5% $CO_2$ for 24 hours was extracted with TriZol® reagent (Invitrogen, MA, USA) according to the manufacturer's instructions. RNA integrity was assessed by electrophoresis on 1% agarose and quantification was performed by absorbance analysis using a NanoDrop™ 2000 spectrophotometer (Thermo Fisher Scientific). cDNAs were prepared from DNase (Promega, WI, USA)-treated total RNA samples (290 ng) using ImProm-II reverse transcriptase (Promega) and oligo-dT. Quantitative real-time PCR (qRT-PCR) was performed on a Fast 7500 real-time PCR system (Applied Biosystems, MA, USA) with the following thermal cycling conditions: 95°C for 10 min followed by 40 cycles of 95°C for 15 s, 55°C for 15 s, and 60°C for 60 s. Platinum® SYBR® green qPCR Supermix (Invitrogen) was used as the reaction mix and supplemented with 5 pmol of each primer (see Data Set S2, Sheet 5, for sequences) and 8 ng of cDNA template for a final volume of 10 µl. All experiments were performed in biological triplicate and each sample was analyzed in triplicate for each primer pair. Melting curve analysis was performed at the end of the reaction to confirm the presence of a single PCR product. Data were normalized to levels of *ACT1*, which was included in each set of PCR experiments. Relative expression was determined using the $2^{-\Delta Ct}$ method.
